# Recapitulating the frataxin activation mechanism in an engineered bacterial cysteine desulfurase supports the architectural switch model

**DOI:** 10.1101/2020.10.06.326603

**Authors:** Shachin Patra, Cheng-Wei Lin, Manas K. Ghosh, Steven M. Havens, Seth A. Cory, David H. Russell, David P. Barondeau

**Author notes:** To whom correspondence should be addressed: Department of Chemistry, Texas A&M University, College Station, TX 77842, USA. **Author Contributions:** S.P., C.L., M.K.G., and S.M.H. performed the research; all authors were involved in the design of the research, analyzing data, and writing the manuscript.

## Abstract

Iron-sulfur (Fe-S) clusters have a key role in many biochemical processes and are essential for most life forms. Despite recent mechanistic advances in understanding the Fe-S cluster biosynthetic pathway, critical questions remain unresolved. Although human NFS1 and *E. coli* IscS share ∼60% sequence identity, NFS1 exhibits low activity and requires activation by the Friedreich’s ataxia protein frataxin (FXN) for *in vivo* function. Surprisingly, structures of the human complex reveal three distinct quaternary structures with one form exhibiting the same subunit interactions as IscS. An architectural switch model has been proposed in which evolutionarily lost interactions between NFS1 subunits results in the formation of low-activity architectures; FXN binding compensates for these lost interactions and facilitates a subunit rearrangement to activate the complex. Here, we used a structure and evolution-guided approach to identify three conserved residues proposed to weaken interactions between NFS1 subunits and transplanted these amino acids into IscS. Compared to native IscS, the engineered variant had a 4000-fold weaker dimer interface and diminished activity that correlated with the absence of the second catalytic subunit. Remarkably, the addition of the FXN homolog to the engineered variant stimulated the decay of the Cys-quinonoid pyridoxal 5’-phosphate intermediate, shifted IscS from the monomeric to dimeric form, and increased the cysteine desulfurase activity, reproducing results from the human system and supporting the architectural switch model. Overall, these studies indicate a weakening of the homodimeric interface was a key development during the evolution of the eukaryotic system and provide new insights into the role of FXN.

## INTRODUCTION

Iron-sulfur (Fe-S) clusters are ubiquitous protein cofactors that play critical roles in a variety of cellular processes such as aerobic respiration and the biosynthesis of DNA, RNA, proteins, metabolites, natural products, and other protein cofactors (1). Over the course of evolution, sophisticated biosynthetic pathways have developed for the assembly and insertion of appropriate Fe-S cluster cofactors for enzymes in these processes (2,3). The best-studied Fe-S biosynthetic pathway is the ISC system, likely due to its wide-spread occurrence in prokaryotes and its critical role in the mitochondrial matrix of eukaryotes (4). Despite the bacterial and mitochondrial ISC pathways having analogous proteins with high sequence homology and similar mechanisms for the formation and distribution of Fe-S cluster cofactors, there are some notable differences. In particular, the eukaryotic Fe-S assembly system appears to have incorporated new components, repurposed the function, and altered enzyme activity profiles as an apparent functional control mechanism for the pathway (5,6). Understanding evolutionary modifications to these systems are therefore critical for elucidating the regulatory and enzymatic mechanisms of Fe-S cluster assembly and for treating human diseases associated with defects in the ISC biosynthetic pathway, including the incurable neurodegenerative disease Friedreich’s Ataxia (FRDA) (7).

The eukaryotic modifications to the ISC system are focused around the central enzyme in the pathway, the pyridoxal-5’-phosphate (PLP) dependent cysteine desulfurase that converts L-cysteine into L-alanine and provides sulfur for Fe-S cluster synthesis on the scaffold protein IscU (8). The *Escherichia coli* IscS is a group I stand-alone homodimeric enzyme with typical cysteine desulfurase activity (*k*_*cat*_ ∼ 7.5 min^-1^) (9-13). In contrast, the human cysteine desulfurase NFS1, which is 60% identical to *E. coli* IscS, is unstable, essentially inactive, and prone to aggregation (14). NFS1 is stabilized by forming a functional complex with the eukaryotic-specific LYRM protein ISD11 (15,16) and the mitochondrial acyl carrier protein (ACP) (5,6,17). This so-called SDA (NF<u>S</u>1 + IS<u>D</u>11 + <u>A</u>CP) complex exhibits very low cysteine desulfurase activity (*k*_*cat*_ ∼ 0.60 min^-1^) (5) but is stimulated by over an order of magnitude by the Friedreich’s ataxia protein frataxin (FXN) in a scaffold protein (ISCU2) dependent manner (18-20). This activity is further stimulated by the presence of iron. Interestingly, both FXN and the *E. coli* homolog CyaY stimulate the cysteine desulfurase activity of the eukaryotic complex but do not impact the cysteine desulfurase activity in the bacterial system (9,21). Overall, these data point to fundamental differences in the bacterial and mitochondrial cysteine desulfurase enzymes that engender distinct regulation for the assembly of Fe-S clusters.

Three different architectures of the mitochondrial cysteine desulfurase complex SDA_ec_ (human NFS1-ISD11 with *E. coli* ACP) were determined; one form exhibits interactions between catalytic subunits similar to IscS, whereas two forms have distinct alternate protein interfaces. The first “open” structure exhibited an α_2_β_2_γ_2_ quaternary structure with ISD11 molecules mediating interactions between two NFS1-ISD11-ACP_ec_ protomers (Fig. 1A). Unlike the extensive dimer interface in the IscS structure (10,11), there are few direct interactions between the NFS1 subunits in this open architecture. As a consequence, the channel that guides the mobile S-transfer loop cysteine to the active site for the intermolecular sulfur transfer reaction is incomplete and the PLP cofactor is solvent-exposed. The second “closed” structure also featured an α_2_β_2_γ_2_ quaternary structure with NFS1-NFS1 interactions more similar to IscS (Fig. 1B), but with a relative rotation of the catalytic subunits that move the PLP cofactors 5 Å closer together and place structural elements from the other subunit in a position to potentially inhibit the trajectory of the mobile S-transfer loop. The third “ready” architecture (Fig. 1C), which has only been observed structurally in the presence of additional proteins ISCU2 (22) or ISCU2 plus FXN (23), also exhibits an α_2_β_2_γ_2_ quaternary structure with NFS1-NFS1 interactions similar to the IscS structure (Fig. 1D) (10,11). These different architectures raised questions as to which of these forms govern the solution behavior of the human cysteine desulfurase and result in the different biochemical properties compared to the analogous prokaryotic enzymes.

**Figure 1.**
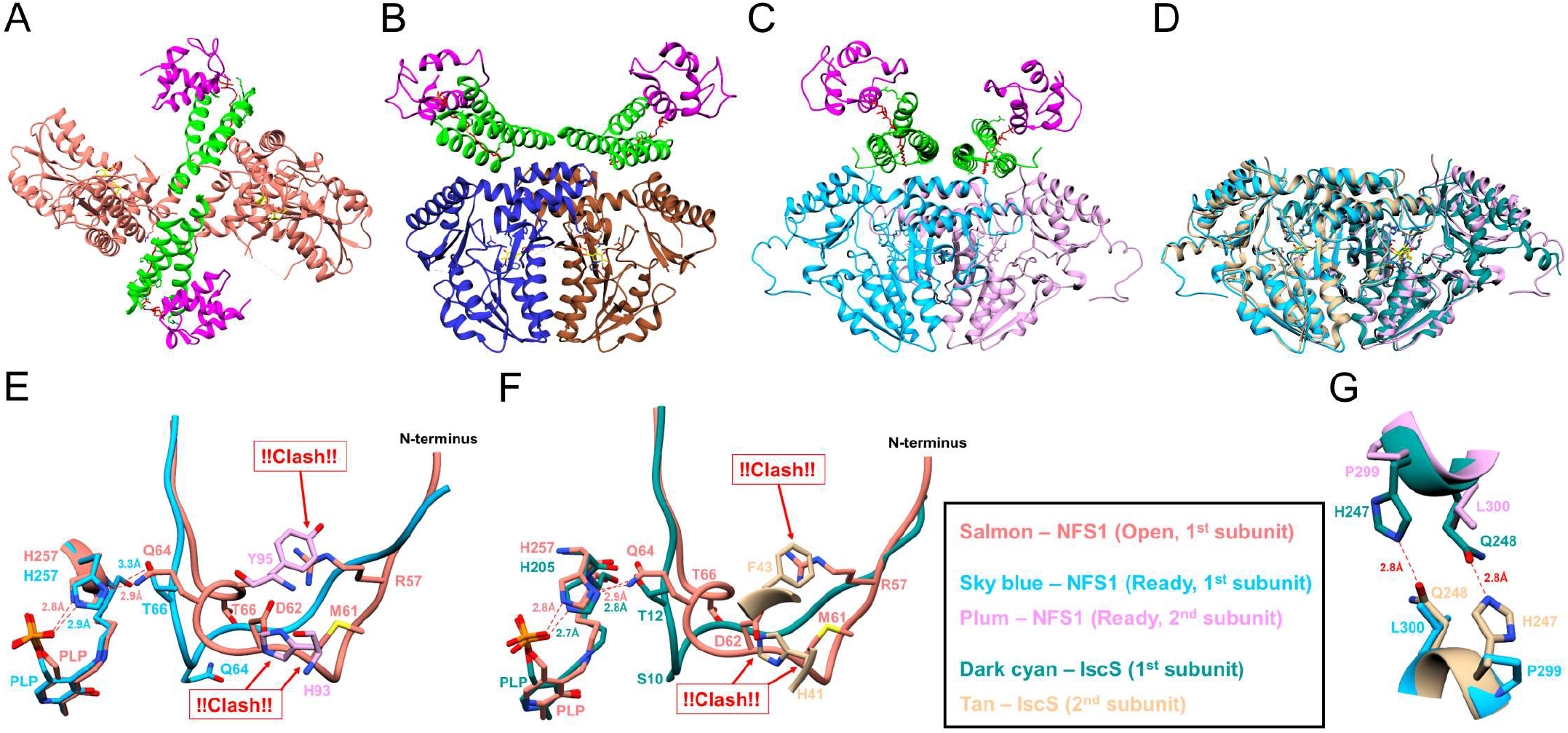
Comparison of the different human cysteine desulfurase architectures and identification of residues that may destabilize interactions between catalytic subunits of the ready form. The human cysteine desulfurase complex (SDA_ec_) in the **(A)** open (pdb code: 5USR), **(B)** closed (pdb code: 5WGB), and **(C)** ready (pdb code: 6NZU) α_2_β_2_γ_2_ architectures. (**D**) Overlay of the *E. coli* IscS (pdb code: 3LVM) with the NFS1 subunits in the ready form. **(E)** Overlay of the region near the PLP cofactor for the open and ready forms of NFS1. The residues Q64 and T66 form hydrogen bonds with H257 in the open and ready forms, respectively. The resulting conformation of this region in the open form sterically occludes the second subunit of NFS1 from forming the ready architecture. (**F**) Overlay of the region near the PLP cofactor for the open form with IscS. Notably, the conformation of the N-terminal region of IscS closely resembles that of NFS1 subunits in the ready form. **(G)** Overlay of IscS with the ready form of NFS1 shows the hydrogen bonds between the IscS H247 Q248 pair with their counterparts from the second subunit. Color scheme: open NFS1 (salmon), closed NFS1 (blue and sierra brown), ready NFS1 (sky blue and plum), ISD11 (green), ACP (magenta), IscS (dark cyan and tan), PLP (yellow), acylated 4’-phosphopantetheine (red).

More recently, stopped-flow and radiolabeling experiments coupled to previous biochemical studies revealed a link between the function of the NFS1 mobile S-transfer loop cysteine and FXN-based activation of the cysteine desulfurase complex (24-26). The addition of FXN was shown to facilitate acid-base, nucleophile, and sulfur transfer functions of the mobile S-transfer loop cysteine in the assembly of Fe-S clusters. Specifically, FXN accelerates the decay of the Cys-quinonoid PLP intermediate, generation of a persulfide species on NFS1, and transfer of sulfur from NFS1 to ISCU2. An architectural switch model has been proposed in which the SDA_ec_ complex exists as low activity species that convert to what we now call the ready form upon FXN binding. The low activity species is composed of the open and/or closed forms, which both exhibit altered substrate binding sites compared to the ready form and lack the structural elements from the second catalytic subunit to guide the mobile S-transfer loop to the active site.

We hypothesized that substitutions were integrated during the evolution of the eukaryotic cysteine desulfurase that weakened the interactions between the catalytic subunits, which led to an increase in monomer concentration. Subsequent interactions between the monomeric form and ISD11/ACP led to the formation of the open architecture. To test this hypothesis, we identified three conserved residues in eukaryotic cysteine desulfurases that may contribute to the weaker NFS1-NFS1 interface for the SDA_ec_ complex and transplanted those residues into *E. coli* IscS. In contrast to native IscS, the resulting S10Q, H247P, Q248L variant (IscS^QPL^) was mostly monomeric at physiologically relevant concentrations and exhibited diminished Cys-quinonoid decay and cysteine desulfurase activities, which were both stimulated by the addition of either CyaY or FXN in a scaffold protein-dependent manner. Importantly, the biochemical properties and activity profile of the IscS^QPL^ variant correlate well with those of the SDA_ec_ complex and suggest that the weakening of the bacterial cysteine desulfurase and monomer formation was a key development during the evolution of the eukaryotic system.

## RESULTS

### Identification of Residues that May Weaken the Interface Between Catalytic Subunits of the Eukaryotic Cysteine Desulfurase

Our initial objectives were to identify amino acid substitutions in the human enzyme that might explain the origin of the multiple SDA_ec_ architectures and provide further insight into the distinct functional properties of the prokaryotic and eukaryotic enzyme complexes. We manually searched for NFS1 residues that are highly conserved in eukaryotes, different than the equivalent residues in IscS, and have the potential to weaken the interface between NFS1 subunits in the SDA_ec_ complex. The search resulted in the identification of two sets of substitutions in the eukaryotic cysteine desulfurases.

The first substitution set replaces *E. coli* IscS residue S10 with NFS1 residue Q64. In some of the NFS1 subunits in the open architecture, Q64 mediates a hydrogen bond to a conserved histidine residue H257 (equivalent to IscS H205) that also hydrogen bonds to the PLP moiety (Figs. 1E and 1F). In contrast, IscS S10 cannot make this interaction, and a conserved threonine (T12 in IscS and T66 in NFS1) from an N-terminal motif in IscS and the closed and ready SDA_ec_ architectures (L59-T66 in NFS1; Fig. S1) makes this interaction instead (Figs. 1E and 1F). The different position in the N-terminal motif of the SDA_ec_ open architecture due to the Q64-H257 hydrogen bond sterically precludes the second NFS1 subunit from generating the ready SDA_ec_ architecture (Fig. 1E). Thus, the highly conserved Q64 (Fig. S2A) allows the N-terminal motif to adopt two conformations: one that allows and one that disfavors an IscS-like ready architecture. Most, if not all, eukaryotic cysteine desulfurases contain glutamine at this position. In the α-proteobacteria clade, which shares ancestry with eukaryotic mitochondria, this position is primarily glutamine, although some IscS homologs have asparagine or histidine residues at this position (Fig. S2B). The glutamine, asparagine, and histidine side-chains share the hydrogen-bonding ability of NFS1 Q64 and may also trigger a dimer-destabilizing structural rearrangement. Conversely, IscS paralogs from β-proteobacteria and γ-proteobacteria, which includes *E. coli*, contain residues at this position with small aliphatic side-chains, such as serine, alanine, or glycine, that are unable to interact with the conserved histidine and would not be expected to facilitate the N-terminal structural rearrangement of the open SDA_ec_ architecture (Figs. S2A and S2B).

The second substitution set replaces *E. coli* IscS H247 and Q248 with NFS1 P299 and L300. IscS H247 and Q248 are at the dimeric interface of IscS and form hydrogen bonds with their counterparts from the second IscS subunit in the dimer (Fig. 1G). The corresponding residues in NFS1 (Fig. S1), P299 and L300, cannot form these hydrogen bonds and should weaken the homodimeric protein-protein interactions of the ready architecture. Phylogenetic analysis revealed that the NFS1 residues are highly conserved among eukaryotes and mostly conserved in α-proteobacteria (Fig. S2A). In contrast, the analogous *E. coli* residues (H247 and Q248) are highly conserved in the β-proteobacteria and γ-proteobacteria clades (Fig. S2C). Overall, three invariant human residues, Q64, P299, and L300, were identified that may contribute to a weaker interface between NFS1 catalytic subunits of the human SDA_ec_ complex.

### The S10Q, H247P, and Q248L Substitutions Favor Monomeric IscS

We tested whether transplanting the three identified eukaryotic residues into the equivalent positions of *E. coli* IscS would weaken the dimer interface and allow the prokaryotic enzyme to recapitulate functional properties of NFS1. Substitutions were incorporated with site-directed mutagenesis and the resulting IscS variants were purified to homogeneity. Size exclusion chromatography (SEC) experiments were used to assess monomeric and dimeric fractions for the IscS variants. At a concentration of 1 μM, native IscS was mostly dimeric, the S10Q variant (IscS^S10Q^) was a roughly equal mixture of monomeric and dimeric species, the H247P Q248L variant (IscS^PL^) had a larger population of monomer than dimer, and the S10Q, H247P, Q248L variant (IscS^QPL^) was mostly monomeric (Fig. 2). SEC analysis of these variants at concentrations ranging from 0.5 μM to 10 μM demonstrated some concentration-dependent effects (Fig. S3). Native IscS was primarily a dimer at all concentrations tested. The IscS^S10Q^ and IscS^PL^ variants exhibited a mixture of monomeric and dimeric peaks at lower concentrations, whereas they existed as mostly dimeric species when the concentration was increased. In contrast, the IscS^QPL^ variant was mostly monomeric and only exhibited roughly equimolar amounts of monomer and dimer when the concentration was increased to ∼50 μM (Figs. S3 and S4).

**Figure 2.**
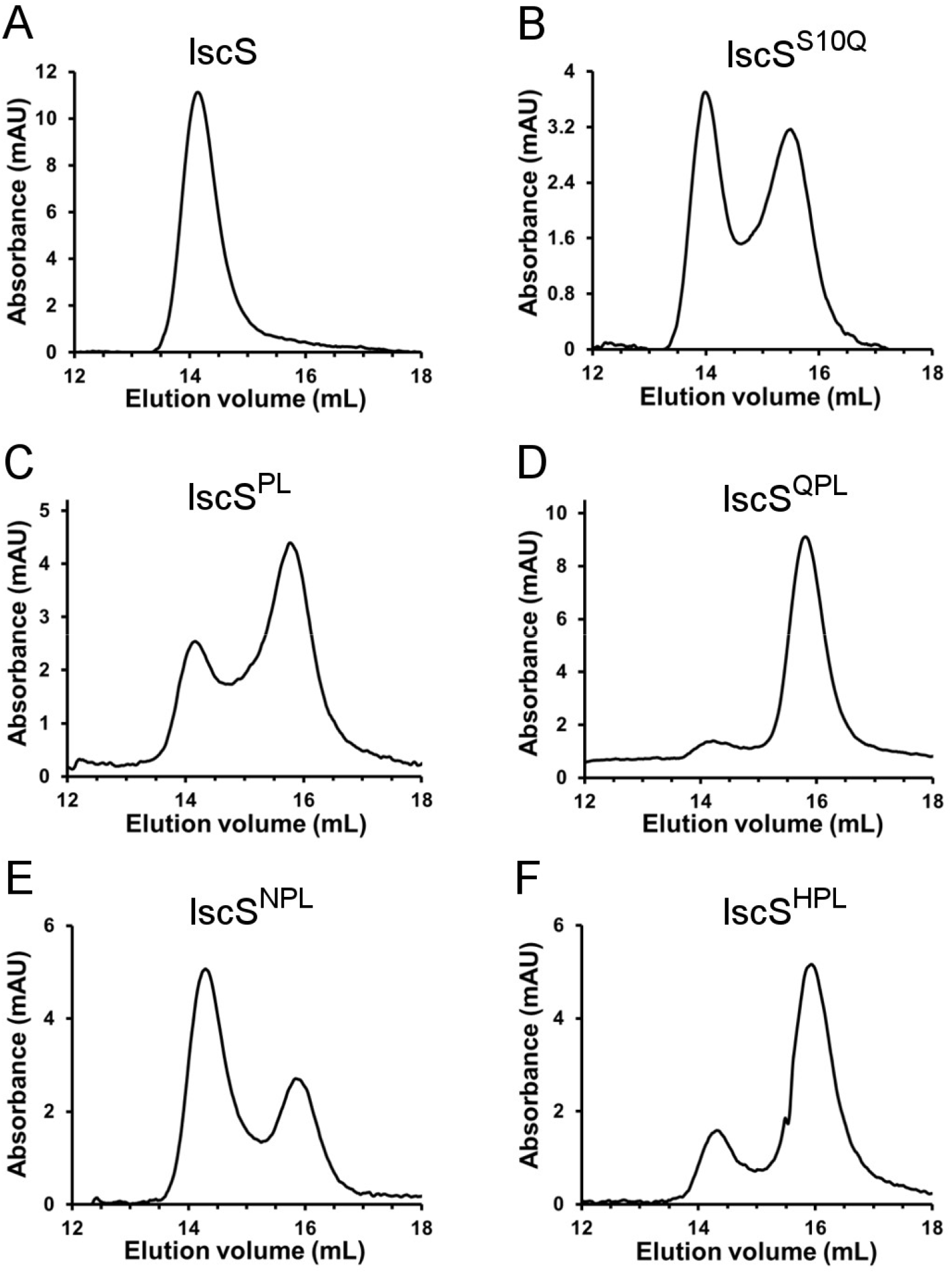
Effect of substitutions on the oligomeric state of IscS. Size-exclusion chromatographic (SEC) analysis of (**A**) native IscS and the (**B**) IscS^S10Q^, (**C**) IscS^PL^, (**D**) IscS^QPL^, (**E**) IscS^NPL^, and (**F**) IscS^HPL^ variants. Proteins were prepared at a concentration of 1 µM in the presence of 2.5 mM TCEP under anaerobic conditions. The peaks with elution volumes of ∼14 and 15.8 mL were estimated to have molecular masses of 91 (dimer) and 45 (monomer) kDa, respectively. SEC analyses at different protein concentrations are shown in Figure S3.

We also tested the impact of the S10N and S10H substitutions on the oligomeric state of the IscS^PL^ variant. These substitutions are found in IscS enzymes from α-proteobacteria lineages (Fig. S2B) and could potentially form hydrogen bonds and promote dimer-destabilizing rearrangements similar to NFS1 Q64. The S10N, H247P, Q248L variant (IscS^NPL^) was predominantly dimeric, whereas the S10H, H247P, Q248L variant (IscS^HPL^) was mostly monomeric but exhibited a larger proportion of dimer than the IscS^QPL^ variant (Figs. 2 and S3). In summary, substitutions at the three positions of IscS all weakened the protein-protein interactions, but the greatest effect was observed for the IscS^QPL^ variant, which mimics the conserved eukaryotic residues.

### The Cysteine Desulfurase Activity of IscS^QPL^ is Stimulated by the Scaffold Proteins, FXN/CyaY, and Fe^2+^

We then sought to determine whether these dimer-weakening substitutions affected the cysteine desulfurase activity of IscS. When assayed under standard conditions, the IscS^S10Q^ (*k*_*cat*_ = 1.9 min^-1^), IscS^PL^ (*k*_*cat*_ = 4.1 min^-1^), IscS^HPL^ (*k*_*cat*_ = 1.5 min^-1^), and IscS^QPL^ (*k*_*cat*_ = 0.22 min^-1^) variants had activities significantly lower than IscS (*k*_*cat*_ = 8.2 min^-1^) (Table S1 and Fig. S5). The activity of the IscS^QPL^ variant was comparable to the human cysteine desulfurase complex SDA_ec_ (*k*_*cat*_ = 0.6 min^-1^) (5). In contrast, the IscS^NPL^ variant had a cysteine desulfurase activity (*k*_*cat*_ = 8.9 min^-1^) similar to native IscS. The IscS^NPL^ and IscS^HPL^ variants were best fit to a Michaelis-Menten equation that included positive cooperativity with a Hill coefficient between 1.4 and 1.6, unlike native IscS or the other variants (Table S1 and Fig. S5). The activities for the different variants were also determined as a function of protein concentration with a constant amount of substrate (Fig. S6). Notably, variants with the highest cysteine desulfurase activities (native IscS plus the IscS^PL^ and IscS^NPL^ variants) exhibited substrate inhibition at higher protein concentrations. Overall, a clear correlation was found between the percentage of dimer from the analytical SEC experiments and the relative cysteine desulfurase activities for the different IscS variants (Fig. 3).

**Figure 3.**
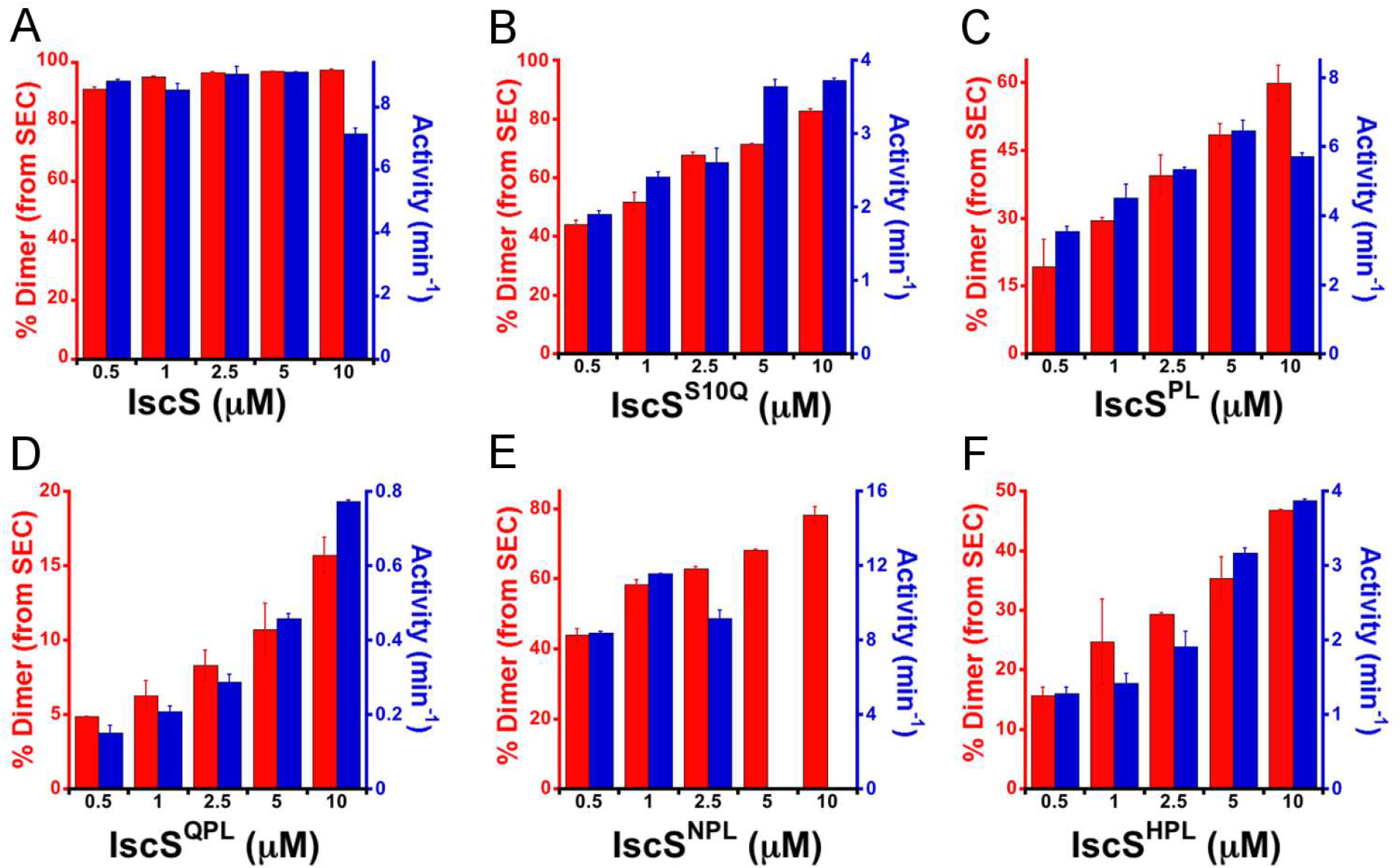
Correlation between the percent dimer and activity for IscS variants. The percent dimer was estimated from SEC data and plotted with the activity of the (**A**) native IscS and the (**B**) IscS^S10Q^, **(C)** IscS^PL^, **(D)** IscS^QPL^, **(E)** IscS^NPL^, and **(F)** IscS^HPL^ variants at different protein concentrations. The activity was determined by dividing the rate of sulfide produced under saturating conditions of L-cysteine (1 mM) by the enzyme concentration. The activity is directly proportional to the percent dimer at the different variant concentrations except at high protein concentrations for the most active variants. In these cases, the substrate becomes a limiting factor in the rate of sulfide generation and increasing the substrate concentration results in an inhibitory effect. Error bars are replicate errors (n=3).

### Substitutions Have Varied Effects on the Dimeric Binding Constant and Dimeric Activity

Under the assumption that only the dimeric IscS was functional, we determined the dimer dissociation constant (*K*_d_) and activity of the dimer (A_s_) from fits of the plots of observed cysteine desulfurase rates vs. enzyme concentration (Fig. S6). Native IscS was determined to have a relatively tight binding affinity between subunits (*K*_d_ = 0.008 µM) and a dimeric activity of 18.7 min^-1^ (Table S1 and Fig. 4). The IscS^PL^ variant had two orders of magnitude lower dimerization affinity (*K*_d_ = 1.4 µM) but very similar dimeric activity (A_s_ = 18.8 min^-1^), suggesting the effect of H247P and Q248L substitutions on activity is primarily due to reduced levels of the dimer.

**Figure 4.**
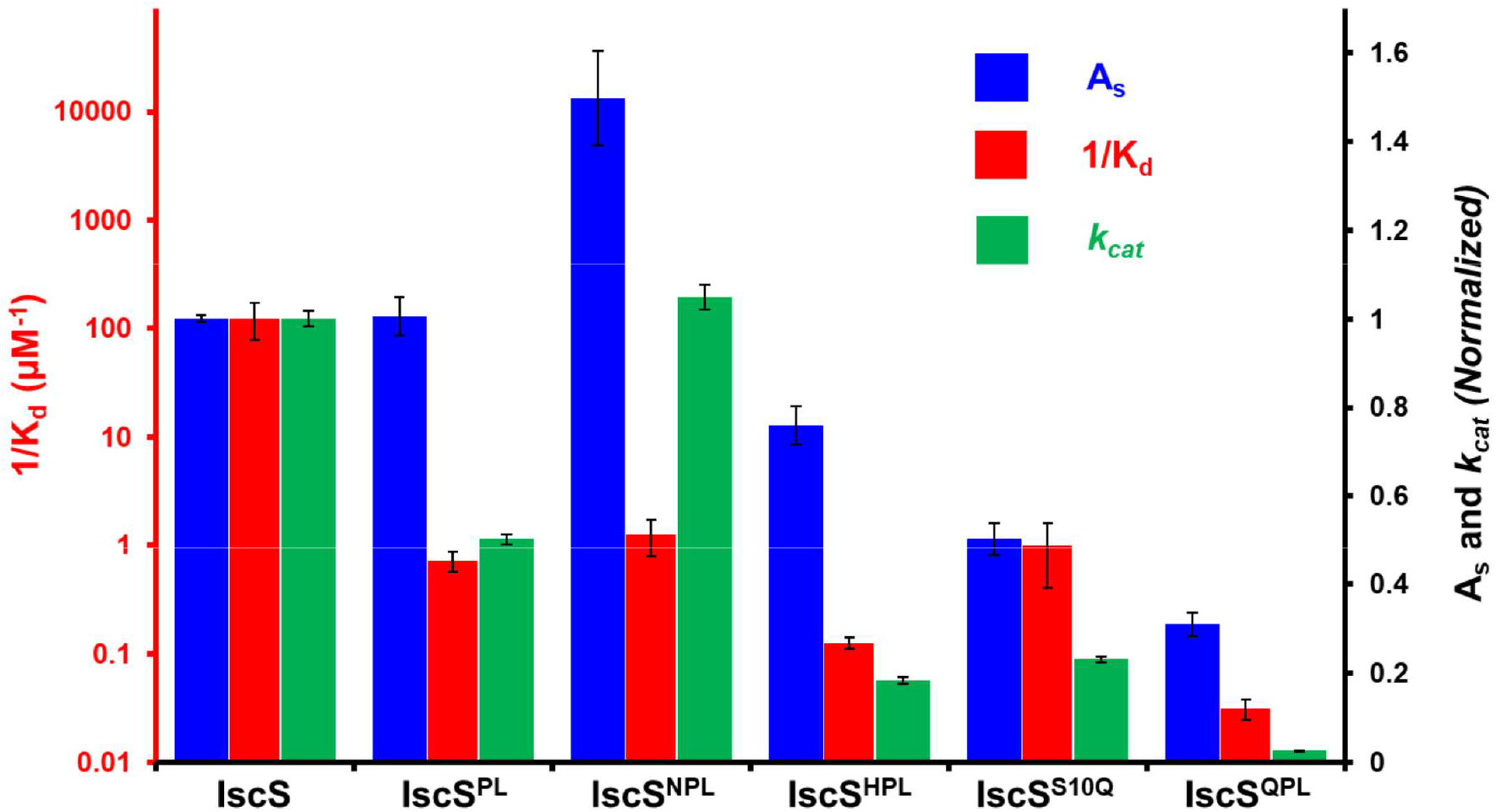
Comparison of the dimeric binding affinity, dimeric activity, and *k*_cat_ for native IscS and IscS variants. The dimeric binding affinity (1/*K*_d_) values are shown on the left Y-axis and the normalized activity (both A_s_ and *k*_*cat*_) values are shown on the right Y-axis for native IscS and the IscS^PL^, IscS^NPL^, IscS^HPL^, IscS^S10Q^, and IscS^QPL^ variants. The *K*_d_, A_s_, and *k*_*cat*_ values are also listed in Table S1. Error bars are replicate errors (n = 3).

Substitutions at position 10 exhibit more complex behavior. The IscS^NPL^ variant had a similar dimeric binding constant (*K*_d_ = 0.8 µM) to the IscS^PL^ variant, indicating that the S10N substitution does not have a large effect on dimerization. However, the dimeric activity for the IscS^NPL^ variant was ∼150% of native IscS, which indicates the S10N substitution enhances the activity and explains the similar *k*_*cat*_ to IscS despite the weaker dimer binding constant (Table S1 and Fig. 4). The IscS^HPL^ variant had an 8 µM dimeric binding constant and 75% of the dimeric activity of the IscS^PL^ variant, indicating that the S10H substitution both weakened the dimer interface and lowered enzymatic activity. Finally, the S10Q substitution further weakened the dimeric binding constant (*K*_d_ = 1 µM for IscS^S10Q^; *K*_d_ = 32 µM for IscS^QPL^) and had the largest negative effect on the dimeric activity (A_s_ = 9.4 min^-1^ for IscS^S10Q^; A_s_ = 5.8 min^-1^ for IscS^QPL^). These fitted dissociation constants accurately predicted the dimer percentages for the variants as measured by analytical SEC (Fig. S7). Overall, the substitutions at H247P and Q248L weakened the dimer interface, S10N enhanced the dimeric activity, and the S10H moderately and the S10Q greatly weakened the subunit interactions and lowered the dimeric activity.

### The Cysteine Desulfurase Activity of IscS^QPL^ is Stimulated by the Scaffold Proteins, FXN/CyaY, and Fe^2+^

The IscS^QPL^ variant is primarily monomeric and has a low cysteine desulfurase activity, which is comparable to that of the SDA_ec_ complex. We therefore tested whether CyaY/FXN could stimulate the activity of the IscS^QPL^ variant similar to the activation of the eukaryotic SDA_ec_ complex (5). Combining the IscS^QPL^ variant with the *E. coli* scaffold protein IscU (S^QPL^U) or the human scaffold protein ISCU2 (S^QPL^U_hs_) resulted in an approximately threefold enhancement of the cysteine desulfurase activity (Figs. 5 and S8). This result was unexpected, as the addition of the scaffold protein to either native IscS or the SDA_ec_ complex results in a slight reduction of cysteine desulfurase activity (9). The addition of either CyaY or FXN resulted in a further rate enhancement in a scaffold protein-dependent manner, similar to the FXN-based activation of the SDA_ec_ complex. The extent of the rate enhancement was dependent on the combination of IscU/ISCU2 and CyaY/FXN, with the *E. coli* protein complex (IscS^QPL^ + IscU + CyaY or S^QPL^UC) exhibiting the maximum rate (Fig. 5). The addition of ferrous iron resulted in a modest additional increase in activity (Fig. S9). Interestingly, the iron-based stimulation and overall ∼10 fold rate enhancement for the S^QPL^UC complex relative to the IscS^QPL^ variant is highly reminiscent of the ∼10 fold FXN-based stimulation for the human system (18).

**Figure 5.**
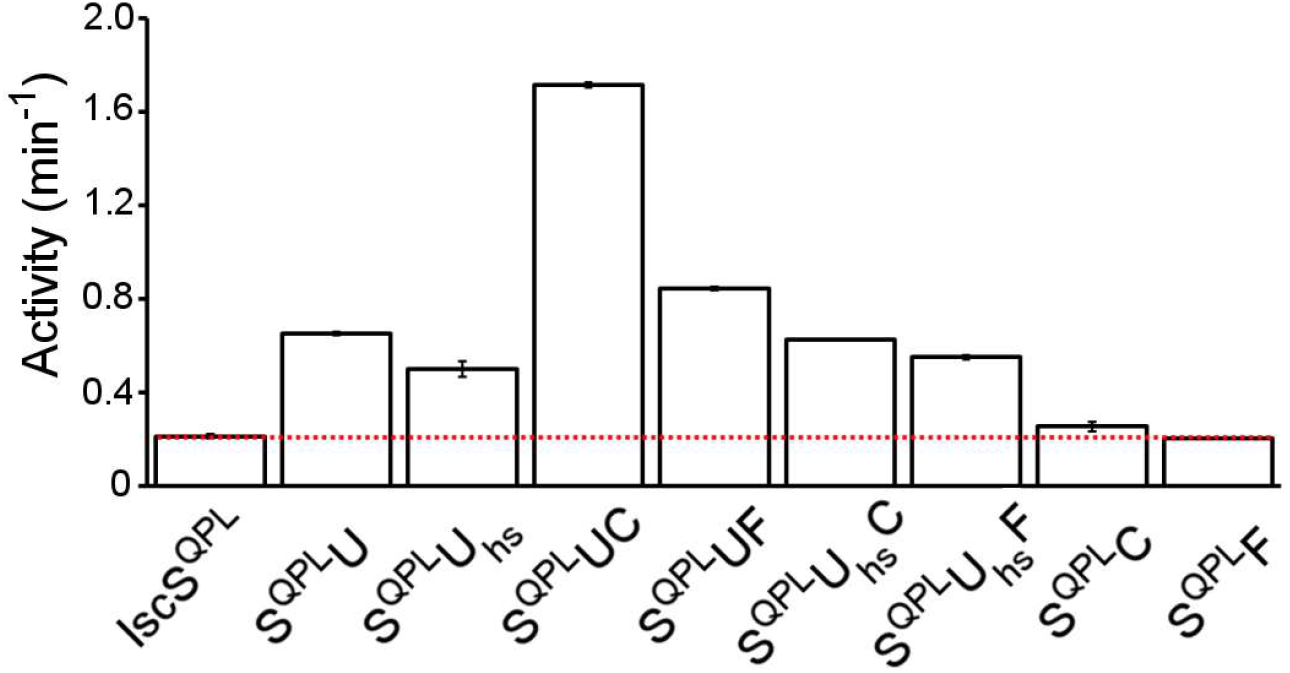
The scaffold protein and CyaY/FXN enhance the cysteine desulfurase activity of the IscS^QPL^ variant. Complexes of the IscS^QPL^ (S^QPL^) variant (0.5 µM) were generated by combining with either 30 µM of the scaffold protein, IscU (U) or ISCU2 (U_hs_), and/or 20 µM of CyaY (C) or FXN (F). Cysteine desulfurase activities were conducted with 1 mM L-cysteine and 4 mM D, L-DTT. The red dotted line is the activity of the IscS^QPL^ variant alone. Error bars represent replicate errors (n=3).

### Both the Scaffold Protein and CyaY/FXN Promote Dimerization to Activate IscS^QPL^

In the architectural switch model, the human cysteine desulfurase complex exists as an equilibrium mixture of species dominated by low activity forms; FXN binding shifts this equilibrium toward the IscS-like dimer of the ready form and positions the NFS1 mobile S-transfer loop to function as an acid, nucleophile, and sulfur-transfer species to activate the complex (5,24). Here, we tested if the activity stimulation of the IscS^QPL^ variant operates through a similar mechanism. First, we used SEC to test if binding IscU shifted the equilibrium of the IscS^QPL^ variant toward the dimeric form. The IscS^QPL^ sample exhibited a major peak associated with the monomer form and a minor peak consistent with the dimeric form (Fig. 6A). The addition of IscU decreased the amount of monomer with a concomitant increase in the dimer. IscU also shifted both the monomeric and dimeric peaks to smaller elution volumes, consistent with IscU binding both monomeric and dimeric IscS. To confirm this, we analyzed both IscS^QPL^ and S^QPL^U samples by native ion mobility mass spectrometry (IM-MS). The native IM-MS data for IscS^QPL^ revealed more monomer than dimer, consistent with the SEC data (Figs. 6B and S10A). In contrast, after adding IscU the total amount of dimeric IscS, including both uncomplexed and IscU-bound species, was greater than the total amount of monomeric IscS (Figs. 6B and S10B). Moreover, monomeric IscS^QPL^ exists as equal populations of IscS^QPL^ and S^QPL^U, whereas most of the dimeric IscS^QPL^ is in a heterotetrameric S^QPL^U assembly (Fig. 6C). Overall, the native MS and SEC results established that both monomeric and dimeric forms of the IscS^QPL^ variant can bind to the scaffold protein but that IscU binding favors the dimeric form.

**Figure 6.**
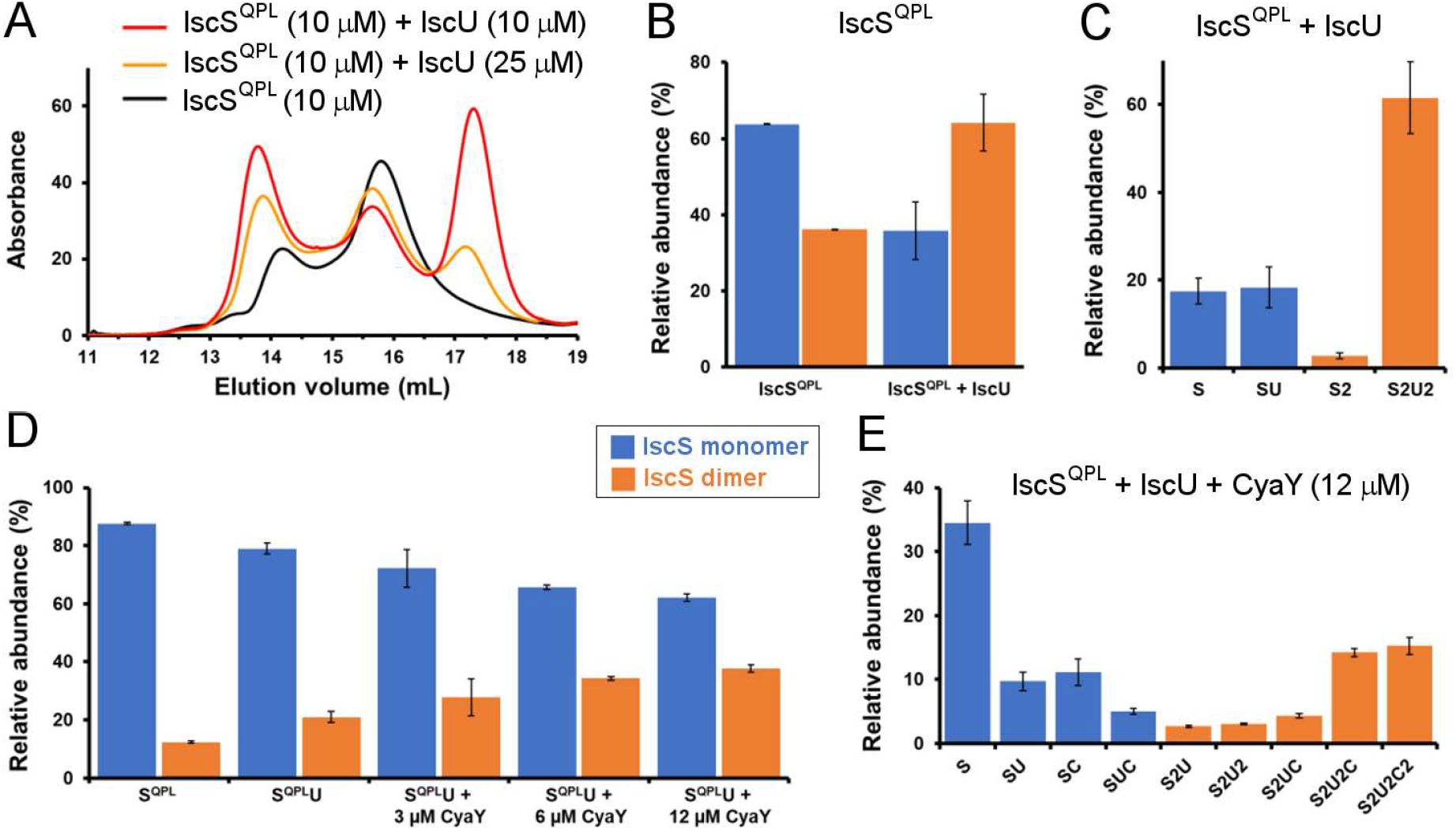
The IscS^QPL^ variant dimerizes in the presence of IscU and CyaY. **(A)** The IscS^QPL^ variant (10 µM) was mixed with IscU (10 or 25 µM) in the presence of 2.5 mM TCEP and analyzed using analytical size exclusion chromatography. The presence of IscU increased the population of the dimer and shifted both the monomer and dimer peaks towards an elution time consistent with larger species. **(B)** Native IM-MS analysis shows the IscS^QPL^ variant (10 µM) exists predominantly as a monomer in the absence of IscU and as a dimer in the presence of IscU (25 µM). **(C)** Detailed analysis of the IscS^QPL^ variant plus the IscU sample in panel B shows that about half of the IscS^QPL^ monomer (S) and almost all the IscS^QPL^ dimer (S2) binds IscU. **(D)** Native IM-MS analysis revealed that CyaY also promoted dimerization. Experimental conditions: 4 µM IscS^QPL^ variant, 6 µM IscU, and 0 – 12 µM CyaY. (**E**) The relative abundance of different species detected by native MS for the highest concentration CyaY sample from panel D. The error bars represent replicate errors (n = 3). Abbreviations: IscS^QPL^ = S; IscU = U; CyaY = C.

We then tested whether CyaY also promotes the dimerization of the IscS^QPL^ variant. Initial experiments revealed that the binding of CyaY to the native IscS-IscU complex is dynamic and not appropriate for SEC analysis. Therefore, native IM-MS was used to monitor the species in solution. The IscS^QPL^ variant exhibited a high monomer-to-dimer ratio that shifted slightly toward the dimeric species upon the addition of a 1.5-fold molar excess of IscU (Fig. 6D). A titration of the S^QPL^U complex with CyaY resulted in a further shift from monomeric to dimeric complexes (Figs. 6D and S10C). Further analysis of the distribution of species reveals CyaY binds to both monomeric (with and without IscU) and dimeric forms of the IscS^QPL^ variant and that the dimeric form is enriched in complexes that include CyaY (Fig. 6E).

### Both IscU and CyaY Accelerate Cys-quinonoid Decay Kinetics

In the human system, the addition of FXN accelerates the decay of the Cys-quinonoid PLP intermediate, likely by positioning the mobile S-transfer loop cysteine to function as a proton donor (24). Therefore, we tested whether there was also a link between the formation of the dimeric form of IscS and the rate of Cys-quinonoid decay using stopped-flow experiments. First, we determined that the rate of quinonoid decay for native IscS and the IscS^S10Q^, IscS^PL^, and IscS^QPL^ variants correlates well with the dimeric binding affinity (IscS > IscS^PL^, IscS^S10Q^ > IscS^QPL^) (Figs. 7A and S11; Table S1). Second, we determined that the quinonoid decay kinetics of native IscS were not affected by the addition of IscU (SU complex), CyaY (SC complex), or both IscU and CyaY (SUC complex) (Figs. 7B and S11). Third, in contrast to native IscS, the quinonoid decay rate for the IscS^QPL^ variant was stimulated by the addition of IscU (S^QPL^U complex) and then further accelerated by the addition of CyaY (S^QPL^UC complex) (Figs. 7C and S11), again correlating with the amount of dimeric enzyme. Overall, we were able to recapitulate key biochemical and functional properties of the human cysteine desulfurase complex in the bacterial system by transplanting conserved residues that weakened the dimer interface of IscS and resulted in an enriched population of monomeric species.

**Figure 7.**
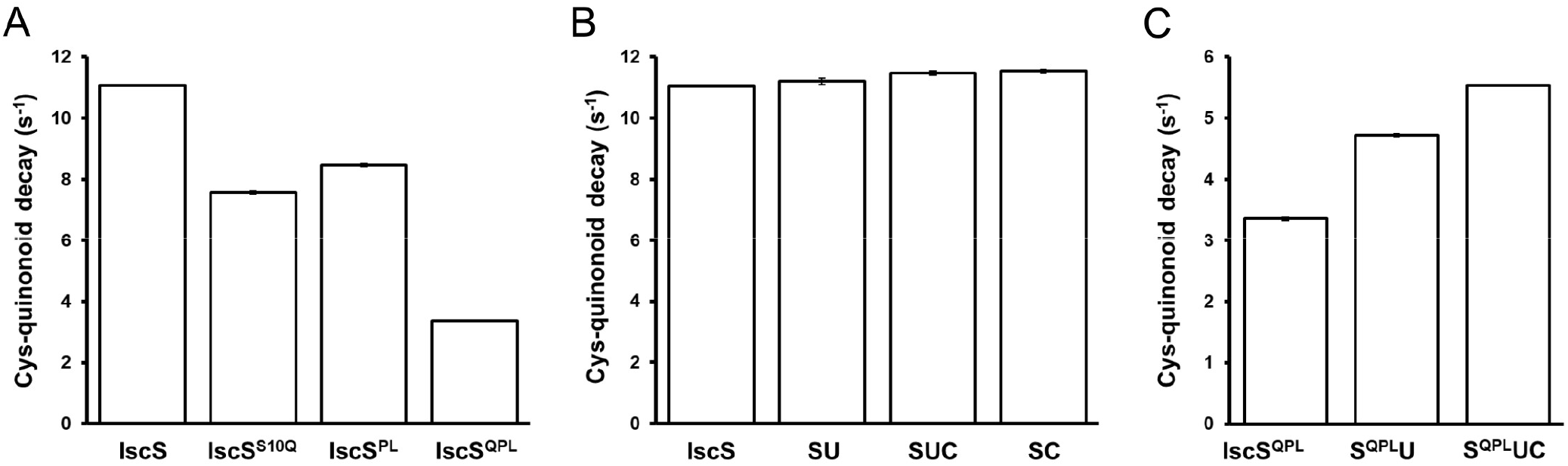
The decay rate of the Cys-quinonoid intermediate is correlated with the amount of dimeric IscS. A comparison of the rates of the Cys-quinonoid decay rates, which were obtained from the fit of the curves in Fig. S11. (**A)** The decrease in the rate of Cys-quinonoid decay positively correlates with the dimer weakening ability of the variants. **(B)** The Cys-quinonoid decay rates for IscS are unaffected by the addition of IscU and CyaY, which do not alter the amount of dimer for native IscS. **(C)** Cys-quinonoid decay of IscS^QPL^ is enhanced by both IscU and CyaY as both increase the dimer proportion of the predominately monomeric IscS^QPL^. Error bars are error in the fit. Abbreviations: IscS = S; IscU = U; CyaY = C.

## DISCUSSION

Although many details of the Fe-S cluster biosynthetic pathway have been elucidated, critical questions remain unresolved that center around differences in the assembly and regulation of the prokaryotic and eukaryotic cysteine desulfurases. Prokaryotic cysteine desulfurases are stable as stand-alone homodimeric enzymes that exhibit similar overall architectures and interactions between subunits (10,27-34). In contrast, structural snapshots of the human cysteine desulfurase complex revealed three distinct quaternary structures that have the same subunit stoichiometry, superimposable NFS1-ISD11-ACP protomers, but use different protein-protein interfaces to generate the open, closed, and ready architectures (Fig. 1) (5,10,22). The biochemical logic for having three different cysteine desulfurase architectures in eukaryotes, and the relative population, interconversion, and physiological functions of the different forms are still poorly understood.

There are also key differences in activity between the prokaryotic and eukaryotic enzymes that are likely linked to the different structural properties. The prokaryotic cysteine desulfurase is a modestly active enzyme; whereas, the eukaryotic complex is a low activity system that requires FXN activation to meet cellular Fe-S cluster biosynthetic demands (5,18,35-37). There is strong evidence that FXN has a role in stimulating the cysteine desulfurase and sulfur transfer chemistry for Fe-S cluster assembly (18-20,23-26,38).

Three competing models have been proposed to explain the origin and relevance of the distinct architectures of the human cysteine desulfurase complex, the differences in activity between the prokaryotic and eukaryotic enzymes in the absence of FXN, and the mechanism of FXN activation. In one proposed model, the Fe-S assembly complex is primarily in the ready architecture both with and without FXN, and the role of FXN is to promote the sulfur transfer reaction from NFS1 to ISCU2 (38,39). However, this model does not take into consideration the open architecture of the SDA_ec_ complex, which has been structurally characterized and shown to be present as a dominant solution component using negative stain electron microscopy (5,22). Moreover, the supporting SAXS and chemical crosslinking studies were not tested for the presence of the open form or appropriately evaluated for their ability to distinguish the closed and ready architectures (22,40,41). This model also fails to explain (*i*) the inherently low cysteine desulfurase activity of the SDA_ec_ complex; (*ii*) the FXN dependent acceleration of both the decay of the Cys-quinonoid intermediate and the formation of the NFS1 persulfide intermediate; and (*iii*) the lack of activity enhancement of IscS by FXN/CyaY (5,18,24).

A second model was proposed based on the cryo-electron microscopy structure of the SDA_ec_UF complex (23) in which FXN induces a conformational change in the mobile S-transfer loop bearing the catalytic cysteine and removes the Zn^2+^ from the ISCU2 active site, thereby activating the eukaryotic cysteine desulfurase (23,42). However, when Zn^2+^ was removed by EDTA, the activity enhancement was modest and to a lower extent compared to the activating effect of FXN (43). Again, this model also neither considers the open architecture, which has been demonstrated to be the dominant SDA_ec_ species in solution, nor explains the inherently low cysteine desulfurase activity of the SDA_ec_ complex (5,18,24). Overall, these models fail to satisfactorily account for all aspects of the activity and regulation for the eukaryotic Fe-S cluster assembly system.

### Architectural switch model

Recently, an architectural switch model has been proposed that can adequately explain the differential activities of the prokaryotic and eukaryotic cysteine desulfurases, the mechanism of FXN-based stimulation, and takes both the open and the ready architectures into account (24). In this model, the eukaryotic cysteine desulfurase complex primarily exists in the low-activity open form in the absence of FXN and is switched to the active ready architecture upon FXN binding. Consistent with this model, a significant population of human NFS1 has been reported to be monomeric (44), which is distinct from *E. coli* IscS that exists almost exclusively as a dimer under physiological conditions (Figs. 2 and S5). Furthermore, this model was also supported by recent studies that show the SDA_ec_ complex exists as an equilibrium mixture of architectures enriched in the open form that can be converted to a single species upon FXN binding (41). We propose that substitutions incorporated during the evolution of the eukaryotic cysteine desulfurase weakened the interactions between the catalytic subunits of the IscS-like ready form and allow the formation of the alternate open SDA_ec_ architecture (Fig. 8). We further propose FXN activates the complex by preferentially stabilizing the ready form to compensate for lost NFS1 subunit interactions that occurred during evolution.

**Figure 8.**
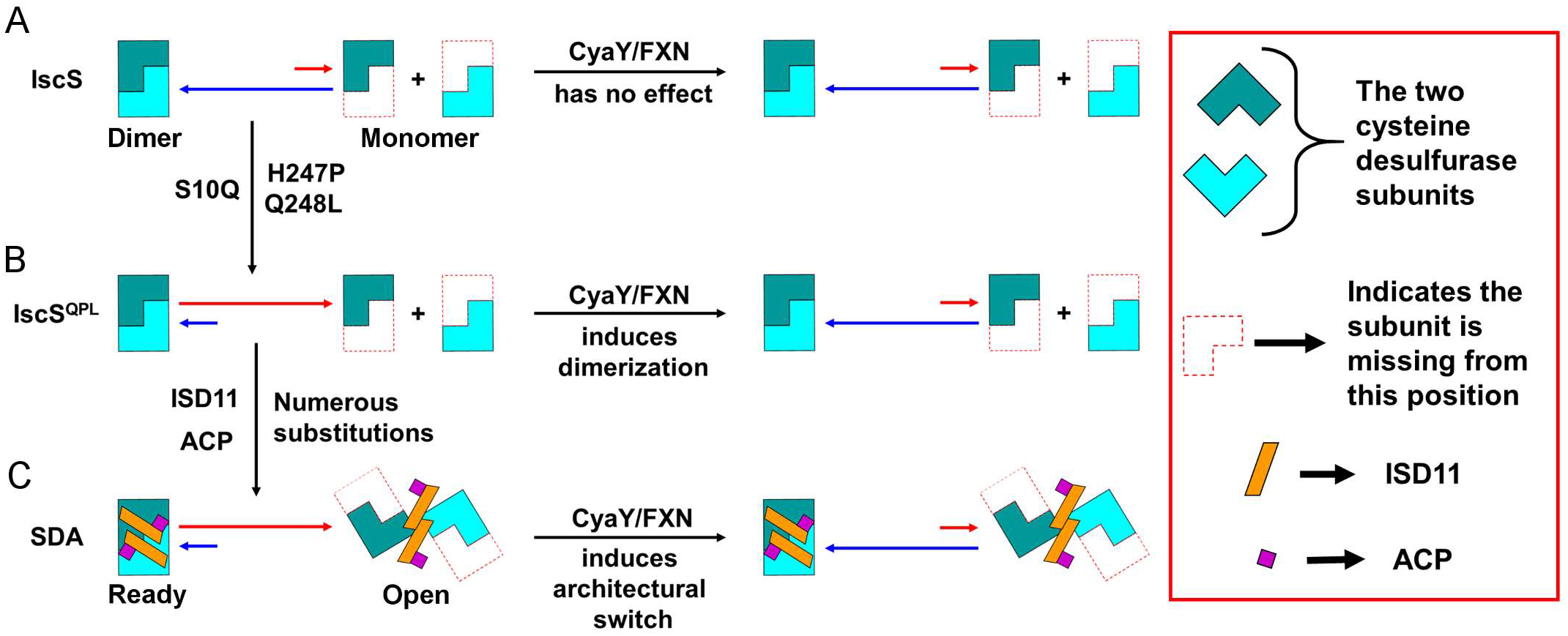
The architectural switch model depicting the mechanism of activation of the eukaryotic cysteine desulfurase by FXN. (**A**) IscS paralogs from the β-proteobacteria and γ-proteobacteria (which includes *E. coli*) exist predominately as a dimer with high activity and are unaffected by the addition of CyaY/FXN. (**B**) Engineering the *E. coli* IscS to incorporate the dimer weakening residues (the Q residue in the N-terminal motif and the HQ pair) that are predominantly found in α-proteobacteria and eukaryotes produced the IscS^QPL^ variant that is primarily monomeric and requires CyaY/FXN to stabilize the high-activity dimeric form and enhance activity. (**C**) During the evolution of eukaryotic NFS1, ISD11 and ACP were incorporated that allowed the generation of the open and ready architectures, which are functionally equivalent to monomeric and dimeric IscS, respectively. Analogous to IscS^QPL^, the SDA complex predominantly exist in the open form, has low activity, and requires CyaY/FXN binding to activate the complex by shifting the equilibrium to the high-activity ready form.

To test this model, a structure and sequence guided approach identified two amino acid motifs that might contribute to weaker interactions between the catalytic subunits of the human cysteine desulfurase complex. Transplanting the human residues from these motifs into *E. coli* IscS generated the IscS^QPL^ variant that exhibited a 4000-fold weaker dimer interface and was primarily monomeric. The IscS^QPL^ variant also exhibited very low cysteine desulfurase activity that was similar to that of the SDA_ec_ complex (5). A strong correlation was also observed between the dimer-to-monomer ratio and the activity for IscS variants (Fig. 3), indicating that dimerization is critical for activity. This is consistent with the proposal that the second subunit, when it occupies the adjacent position to the first subunit of the IscS-like ready SDA_ec_ architecture, guides the catalytic cysteine on the mobile S-transfer loop towards the active site PLP cofactor and enhances all of the chemical steps involving the catalytic cysteine that lead to high cysteine desulfurase activity (24).

Consistently, we also observe a correlation between the amount of dimeric IscS and the rate of decay of the Cys-quinonoid intermediate (Fig. 7A), which is dependent on the ability of the mobile S-transfer loop to function as a proton donor (45-47). Moreover, CyaY induced dimerization of the IscS^QPL^ variant accelerated the decay of the Cys-quinonoid intermediate and induced a ∼10-fold stimulation of the cysteine desulfurase activity. These results for the IscS^QPL^ variant closely mimic the effects upon FXN binding to the SDA_ec_U complex (9,18,24) and the proposed subunit rearrangement and stabilization of the ready architecture (24,41). Taken together, these results show that the open and ready forms of the SDA_ec_ complex are functionally equivalent to the monomeric and dimeric IscS, respectively (Fig. 8). Both the ready form and dimeric IscS have a properly positioned second subunit that facilitates faster Cys-quinonoid decay kinetics, persulfide formation on the catalytic cysteine, and sulfur transfer to the scaffold protein. In contrast, both the open architecture and the monomeric IscS lack a properly positioned second subunit and have decreased cysteine desulfurase activity and slower Cys-quinonoid decay kinetics.

### Evolution of a weaker interface between cysteine desulfurase catalytic subunits

This work also points towards a likely scenario for the evolution of the eukaryotic cysteine desulfurase and the FXN-based activation mechanism. Analysis of the prevalence of the dimer-weakening residues for IscS in proteobacteria (Fig. S2) suggests that the last common ancestor of the α-, β-, and γ-proteobacteria likely contained a non-Q residue in its N-terminal motif (position 10 in *E. coli* IscS) and a pair of non-HQ residues at the homodimeric interface (Figs. S2 and S12). During evolution, the β-proteobacteria and γ-proteobacteria acquired the dimer-stabilizing HQ motif. These additional hydrogen bonding interactions at the dimer interface likely increased the affinity between the IscS subunits and increased the proportion of the active dimeric form. As aerobic conditions inhibit Fe-S cluster assembly and degrade Fe-S clusters, an increase in cysteine desulfurase activity as a result of increased dimer fraction may have been a critical adaptation to the aerobic lifestyle after the great oxidation event (48). A stabilized dimeric form of IscS would not require CyaY to further increase the dimeric fraction, consistent with the lack of CyaY stimulation for native *E. coli* IscS (9), which suggests that CyaY either has lost its function or has some other role in these organisms.

In contrast, most of the α-proteobacteria, especially the Rickettsiales order (49), acquired the dimer-destabilizing glutamine residue, which in NFS1 allows a structural rearrangement of the N-terminus that disfavors the ready form. We propose that this substitution, in combination with the lack of the dimer-stabilizing HQ hydrogen-bonding pair, would have significantly weakened the homodimeric interface resulting in a high degree of monomer and low inherent activity of the enzyme, similar to the IscS^QPL^ variant (Fig. 8). This would have posed a challenge to take advantage of the increased oxygen level and adopt an aerobic lifestyle after the great oxidation event (48). The presence of CyaY, which can enhance the activity of the cysteine desulfurase monomer by facilitating dimerization, might have been critical in overcoming this deleterious situation and meet the cellular biosynthetic demands.

We hypothesize, after the evolution of mitochondria, eukaryotes acquired ISD11, which with the help of the already present ACP could form a complex with NFS1. The NFS1 monomers predominantly gave rise to the functionally equivalent open architecture along with maybe a small fraction of the closed architecture, whereas the NFS1 dimers resulted in the functionally equivalent ready architecture. We further propose that the incorporation of ISD11 and ACP did not significantly affect the monomer-dimer equilibrium of NFS1 (Fig. 8). Consequently, the low-activity open architecture was the predominant species in the absence of FXN. FXN was also able to shift the equilibrium to the high-activity ready architecture resulting in activity enhancement. This is consistent with FXN being a critical component of the eukaryotic Fe-S cluster assembly machinery under aerobic growth conditions but being dispensable at lower oxygen levels (37). Interesting future studies include evaluating whether IscS enzymes in the α-proteobacteria clade do indeed have weaker homodimeric interfaces and functional dependence on CyaY under aerobic conditions.

In summary, this study identified three highly conserved residues in eukaryotic NFS1 that when transplanted into *E. coli* IscS resulted in a 4000-fold weaker dimer interface. The resultant IscS^QPL^ variant was predominantly monomeric at physiological concentration, had similar biochemical properties to that of SDA_ec_, and reproduced the FXN activation phenomena of the human system (9,18,24). A link between the quaternary structure and the mobile S-transfer loop trajectory explains the different Cys-quinonoid decay kinetics and cysteine desulfurase activities and the correlation of the SDA_ec_ ready form with dimeric IscS and the SDA_ec_ open form with monomeric IscS. Our results also provided a likely scenario for the origin of the eukaryotic cysteine desulfurase complex, its low activity, and the activity enhancement by FXN. Overall, these results suggest that the weakening of the homodimeric interface was a key development during the evolution of the eukaryotic system that further support the architectural switch model (24,41), and provide additional insight into the role of FXN with implications in potential therapeutics for the incurable neurodegenerative disease Friedreich’s Ataxia.

## MATERIALS AND METHODS

### Protein Expression and Purification

The SDA_ec_ complex (5), ISCU2 (9), FXN (9), IscS (9), IscU (9), and CyaY (9) were expressed and purified as previously described. The Quick-change protocol (Agilent) was used to introduce point mutations. IscS variants were purified similarly to the native enzyme. Protein concentrations for the SDA_ec_ complex, IscS, and IscS variants were estimated using an extinction coefficient of 6.6 mM^-1^cm^-1^ at 388 nm (in 0.1 M NaOH) (50). Protein concentrations for IscU, ISCU2, CyaY and FXN were estimated in Buffer A (50 mM HEPES, 250 mM NaCl, pH 7.5) using extinction coefficients at 280 nm of 11460 M^-1^cm^-1^, 8490 M^-1^cm^-1^, 28990 M^-1^cm^-1^ and 26030 M^-1^cm^-1^, respectively. Unless otherwise stated, all reactions were carried out in an anaerobic glove box (MBRAUN; maintained at ∼14 °C with O_2_ < 1 ppm).

### Determination of Diversity of Residues Across Different Proteobacterial Lineages

To identify the diversity of residues equivalent to positions 10, 247, and 248 of *E. coli* IscS, homologous sequences were manually curated for each genus within the phylum proteobacteria. The homology decreased as the phylogenetic distance increased making it difficult to classify the sequence as IscS or NifS, the cysteine desulfurase from the NIF Fe-S biosynthetic pathway. Therefore, we used the highly conserved IscS GGG motif (position 232-234 of E. coli IscS) compared to the NifS GGHQ sequence as a marker. We analyzed 60 α-proteobacterial, 103 β-proteobacterial, and 189 γ-proteobacterial IscS sequences. The frequency of a residue at a given position was calculated by dividing the number of sequences with the residue at the given position by the total number of sequences in that lineage. The sequence logos were generated using WebLogo (51) after aligning the protein sequences using Clustal Omega (52).

### Analytical Size-Exclusion Chromatography

IscS variants were diluted in Buffer A to final concentrations of 0.5, 1.0, 2.5, 5.0, and 10 µM in the presence of 2.5 mM TCEP (at pH 8.0). 500 µL of each sample was individually injected into an S200 column (Superdex 200 10/300 GL, GE Healthcare Life Sciences) equilibrated in Buffer A and eluted with a flow rate of 0.5 mL/min. A standard curve was generated using thyroglobulin (669 kDa), apo-ferritin (443 kDa), β-Amylase (200 kDa), alcohol dehydrogenase (150 kDa), albumin (66 kDa), carbonic anhydrase (29 kDa) as molecular weight standards and blue dextran (∼2000 kDa) for determining the void volume (Fig. S13). K_av_ was calculated using the formula K_av_ = (V_e_ – V_v_)/(V_c_ – V_v_), where V_e_, V_v_ and V_c_ are elution volume, void volume, and column volume, respectively. A standard curve was generated by plotting the K_av_ values for the different standards against the log of the molecular weights and the resulting data was fit with a linear equation (y = mx + c). The percent monomer was determined by dividing the estimated area under the monomer peak (elution volume of ∼15.4 to 17 mL) with the total area (monomer + dimer) using UNICORN software (default software of AKTA FPLC, GE Healthcare Life Science), followed by multiplication by 100.

### Cysteine Desulfurase Activity Measurements

Cysteine desulfurase activities were measured for each complex using a slightly modified methylene blue assay (18,53). The 800 µL reaction mixture contained 0.5 µM of native IscS or IscS variants, 4 mM D, L-DTT (final concentration) in Buffer A. The reaction mixture was first incubated for 15 min anaerobically on a heating block at 37 °C. Different concentrations of L-cysteine (0 – 800 µM) were then added, incubated at 37 °C for 6 min and quenched with 20 mM *N, N’*-diphenyl-*p*-phenylenediamine (DPD) (in 7.2 M HCl) and 30 mM FeCl_3_ (in 1.2 M HCl). The samples were incubated for another 20 min, centrifuged at 10000g for 5 min and the absorbance at 670 nm was measured. The amount of sulfide produced was determined using a standard curve made from known amounts of sulfide. Rates ([HS^-^]/([cysteine desulfurase]*min)) were plotted against cysteine concentration and fitted to the Michaelis-Menten equation with cooperativity (below) using Origin software (OriginLab) to obtain the *k*_*cat*_, K_M,_ and Hill coefficient (n). Hill coefficients with n > 1, n < 1 and n = 1 implies positive, negative and no cooperativity, respectively. The Michaelis-Menten equation that includes cooperativity is *y* = (*k*_*cat*_ × *x*^*n*^)/(*K*_M_^*n*^ + *x*^*n*^), where y is the activity, x is the substrate concentration, and n is the Hill coefficient.

### Determination of Dimer Dissociation Constants and Dimeric Activities

The dimer dissociation constants and dimeric activities were calculated based on a fit of the activity at different protein concentrations (54). Cysteine desulfurase activities were determined at final concentrations of 0.5, 1, 2.5, 5, and 10 µM for the IscS variants in the presence of 4 mM D, L-DTT, and 1 mM L-cysteine using the methylene blue assay. We were unable to determine the rate at higher protein concentrations for native IscS and the most active variants as under those conditions increasing the L-cysteine or D, L-DTT concentrations inhibited the reaction. Samples with A_670_ > 1 were diluted 10-fold with a blank solution, 800 µL assay buffer + 100 µL of 20 mM *N,N’*-diphenyl-*p*-phenylenediamine (DPD) (in 7.2 M HCl) + 100 µL of 30 mM FeCl_3_ (in 1.2 M HCl), and remeasured. The equilibrium between the dimer (D) and monomer (M) (D ⇄ 2M) leads to the expression of the dimer dissociation constant 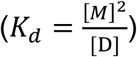, and the total amount of enzyme (*E_t_* = 2[*D*] + [*M*]). Substitution leads to monomer, 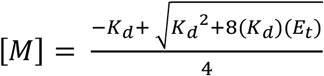, and dimer, 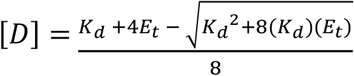, expressions. An initial assumption was made that the monomeric form has no activity and all the activity is due to the dimeric species. Therefore, the observed rate can be described as A_s_ × [D], where A_s_ the dimeric activity (i.e., the activity of the enzyme when present as 100% dimer) and an expression for the observed rate can be generated, 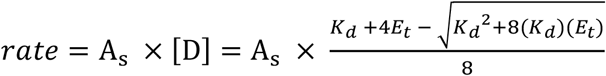; the observed rates were plotted against total enzyme concentration (E_t_) and fit using Origin software (OriginLab) to generate the dimer dissociation constant (*K*_d_) as well as the dimeric activity (A_s_). The binding constant and the above monomer and dimer concentration expressions were used to calculate the amount of monomer and dimer at a given protein concentration.

### Native Ion Mobility Mass Spectrometry (IM-MS)

Native IM-MS experiments were performed on a Synapt G2 instrument (Waters Corporation, U.K.) equipped with an 8k RF generator for ion mobility measurements. Nano-electrospray ionization was performed using a Pt wire inserted into borosilicate tips prepared in house (Sutter 1000) (55). Fresh protein samples including IscS, IscU, and CyaY were buffer exchanged into 200 mM ammonium acetate (pH = 7.5) using Micro Bio-Spin 6 Columns (Bio-Rad). Instrument parameters were tuned to maximize transmission but simultaneously preserve the native-like state of proteins as determined by ion mobility. The instrument was set to a capillary voltage of 1.1 kV, source temperature (30 °C), sampling cone voltage of 20 V, extraction cone voltage of 1 V, trap and transfer collision energy off, and backing pressure (5.07 mbar), trap flow rate at 8 ml/min (3.71 × 10^−2^ mbar), He cell flow rate at 120 ml/min (1.36 × 10^3^ mbar), IMS flow rate at 60 ml/min (2.25 mbar). The T-wave settings for trap (310 ms^-1^/6.0 V), IMS (300 ms^-1^/9-12 V), transfer (65 ms^-^1/2 V), and trap bias (30.0 V). Recorded spectra were deconvoluted MassLynx 4.1 (Waters) and DriftScope v2.1 (Waters). Mass spectra were calibrated externally using a sodium iodide solution and displayed after using the mean smooth method (20 smooth windows, 5 smooths). Quantitation of monomer/dimer intensity was done by extracting corresponding IMS species using DriftScope.

### Stopped-Flow Kinetics for the Cysteine Desulfurase Reaction

Individual 30 µM samples (final concentration) of the native IscS with and without IscU (30 μM; SU complex), CyaY (30 μM; SC complex), and both IscU and CyaY (30 μM of each; SUC complex), IscS variants (IscS^S10Q^, IscS^PL^, and IscS^QPL^), and the IscS^QPL^ variant with IscU (30 μM; S^QPL^U complex), and IscU plus CyaY (30 μM each, S^QPL^UC complex) in assay buffer (50 mM HEPES, 250 mM NaCl, pH 7.5) were combined with 5 mM L-cysteine (final concentration) using a stopped-flow apparatus (KinTek Corporation). Formation and decay of the quinonoid intermediates were followed by monitoring changes in absorbance at 508 nm (24). Traces were fitted with Origin software (OriginLab) to the equation [y = y_0_ + (*k*_1_*[A]_0_/(*k*_2_-*k*_1_))*(exp(-*k*_1_*t)-exp(-*k*_2_*t))], where *k*_1_ and *k*_2_ are rate constants of the formation and decay of intermediates, respectively.

## ACKNOWLEDGMENTS

We wish to thank Dr. Chris Putnam for helpful comments on the manuscript and Professor Tadhg Begley for use of his stopped-flow instrument. Support for this work was funded by the NIH grants R01GM096100 (D.P.B.), R01GM121751 (D.H.R.), and P41GM128577 (D.H.R.) plus the Robert A. Welch grant A-1647 (D.P.B).

## REFERENCES

1. Johnson, D. C., Dean, D. R., Smith, A. D., and Johnson, M. K. (2005) Structure, function, and formation of biological iron-sulfur clusters. Annu Rev Biochem 74, 247–281

2. Ayala-Castro, C., Saini, A., and Outten, F. W. (2008) Fe-S cluster assembly pathways in bacteria. Microbiol Mol Biol Rev 72, 110–125, table of contents

3. Lill, R., and Muhlenhoff, U. (2006) Iron-sulfur protein biogenesis in eukaryotes: components and mechanisms. Annu Rev Cell Dev Biol 22, 457–486

4. Muhlenhoff, U., and Lill, R. (2000) Biogenesis of iron-sulfur proteins in eukaryotes: a novel task of mitochondria that is inherited from bacteria. Biochim Biophys Acta 1459, 370–382

5. Cory, S. A., Van Vranken, J. G., Brignole, E. J., Patra, S., Winge, D. R., Drennan, C. L., Rutter, J., and Barondeau, D. P. (2017) Structure of human Fe-S assembly subcomplex reveals unexpected cysteine desulfurase architecture and acyl-ACP-ISD11 interactions. Proc Natl Acad Sci U S A 114, E5325–E5334

6. Van Vranken, J. G., Jeong, M. Y., Wei, P., Chen, Y. C., Gygi, S. P., Winge, D. R., and Rutter, J. (2016) The mitochondrial acyl carrier protein (ACP) coordinates mitochondrial fatty acid synthesis with iron sulfur cluster biogenesis. Elife 5

7. Rouault, T. A., and Tong, W. H. (2008) Iron-sulfur cluster biogenesis and human disease. Trends Genet 24, 398–407

8. Muhlenhoff, U., Balk, J., Richhardt, N., Kaiser, J. T., Sipos, K., Kispal, G., and Lill, R. (2004) Functional characterization of the eukaryotic cysteine desulfurase Nfs1p from Saccharomyces cerevisiae. Journal of Biological Chemistry 279, 36906–36915

9. Bridwell-Rabb, J., Iannuzzi, C., Pastore, A., and Barondeau, D. P. (2012) Effector role reversal during evolution: the case of frataxin in Fe-S cluster biosynthesis. Biochemistry 51, 2506–2514

10. Cupp-Vickery, J. R., Urbina, H., and Vickery, L. E. (2003) Crystal structure of IscS, a cysteine desulfurase from Escherichia coli. J Mol Biol 330, 1049–1059

11. Shi, R., Proteau, A., Villarroya, M., Moukadiri, I., Zhang, L. H., Trempe, J. F., Matte, A., Armengod, M. E., and Cygler, M. (2010) Structural Basis for Fe-S Cluster Assembly and tRNA Thiolation Mediated by IscS Protein-Protein Interactions. Plos Biol 8

12. Mihara, H., Kurihara, T., Yoshimura, T., Soda, K., and Esaki, N. (1997) Cysteine sulfinate desulfinase, a NIFS-like protein of Escherichia coli with selenocysteine lyase and cysteine desulfurase activities. Gene cloning, purification, and characterization of a novel pyridoxal enzyme. J Biol Chem 272, 22417–22424

13. Mihara, H., and Esaki, N. (2002) Bacterial cysteine desulfurases: their function and mechanisms. Appl Microbiol Biotechnol 60, 12–23

14. Marelja, Z., Stocklein, W., Nimtz, M., and Leimkuhler, S. (2008) A novel role for human Nfs1 in the cytoplasm: Nfs1 acts as a sulfur donor for MOCS3, a protein involved in molybdenum cofactor biosynthesis. J Biol Chem 283, 25178–25185

15. Adam, A. C., Bornhovd, C., Prokisch, H., Neupert, W., and Hell, K. (2006) The Nfs1 interacting protein Isd11 has an essential role in Fe/S cluster biogenesis in mitochondria. Embo J 25, 174–183

16. Wiedemann, N., Urzica, E., Guiard, B., Muller, H., Lohaus, C., Meyer, H. E., Ryan, M. T., Meisinger, C., Muhlenhoff, U., Lill, R., and Pfanner, N. (2006) Essential role of Isd11 in mitochondrial iron-sulfur cluster synthesis on Isu scaffold proteins. Embo J 25, 184–195

17. Cai, K., Frederick, R. O., Tonelli, M., and Markley, J. L. (2017) Mitochondrial Cysteine Desulfurase and ISD11 Coexpressed in Escherichia coli Yield Complex Containing Acyl Carrier Protein. ACS chemical biology 12, 918–921

18. Tsai, C. L., and Barondeau, D. P. (2010) Human frataxin is an allosteric switch that activates the Fe-S cluster biosynthetic complex. Biochemistry 49, 9132–9139

19. Colin, F., Martelli, A., Clemancey, M., Latour, J. M., Gambarelli, S., Zeppieri, L., Birck, C., Page, A., Puccio, H., and Ollagnier de Choudens, S. (2013) Mammalian frataxin controls sulfur production and iron entry during de novo Fe4S4 cluster assembly. J Am Chem Soc 135, 733–740

20. Pandey, A., Gordon, D. M., Pain, J., Stemmler, T. L., Dancis, A., and Pain, D. (2013) Frataxin directly stimulates mitochondrial cysteine desulfurase by exposing substrate-binding sites, and a mutant Fe-S cluster scaffold protein with frataxin-bypassing ability acts similarly. J Biol Chem 288, 36773–36786

21. Adinolfi, S., Iannuzzi, C., Prischi, F., Pastore, C., Iametti, S., Martin, S. R., Bonomi, F., and Pastore, A. (2009) Bacterial frataxin CyaY is the gatekeeper of iron-sulfur cluster formation catalyzed by IscS. Nat Struct Mol Biol 16, 390–396

22. Boniecki, M. T., Freibert, S. A., Muhlenhoff, U., Lill, R., and Cygler, M. (2017) Structure and functional dynamics of the mitochondrial Fe/S cluster synthesis complex. Nat Commun 8, p1287

23. Fox, N. G., Yu, X., Feng, X., Bailey, H. J., Martelli, A., Nabhan, J. F., Strain-Damerell, C., Bulawa, C., Yue, W. W., and Han, S. (2019) Structure of the human frataxin-bound iron-sulfur cluster assembly complex provides insight into its activation mechanism. Nat Commun 10, p2210

24. Patra, S., and Barondeau, D. P. (2019) Mechanism of activation of the human cysteine desulfurase complex by frataxin. Proc Natl Acad Sci U S A 116, 19421–19430

25. Bridwell-Rabb, J., Fox, N. G., Tsai, C. L., Winn, A. M., and Barondeau, D. P. (2014) Human frataxin activates Fe-S cluster biosynthesis by facilitating sulfur transfer chemistry. Biochemistry 53, 4904–4913

26. Parent, A., Elduque, X., Cornu, D., Belot, L., Le Caer, J. P., Grandas, A., Toledano, M. B., and D’Autreaux, B. (2015) Mammalian frataxin directly enhances sulfur transfer of NFS1 persulfide to both ISCU and free thiols. Nat Commun 6, p5686

27. Tirupati, B., Vey, J. L., Drennan, C. L., and Bollinger, J. M., Jr. (2004) Kinetic and structural characterization of Slr0077/SufS, the essential cysteine desulfurase from Synechocystis sp. PCC 6803. Biochemistry 43, 12210–12219

28. Nakamura, R., Hikita, M., Ogawa, S., Takahashi, Y., and Fujishiro, T. (2020) Snapshots of PLP-substrate and PLP-product external aldimines as intermediates in two types of cysteine desulfurase enzymes. FEBS J 287, 1138–1154

29. Blauenburg, B., Mielcarek, A., Altegoer, F., Fage, C. D., Linne, U., Bange, G., and Marahiel, M. A. (2016) Crystal Structure of Bacillus subtilis Cysteine Desulfurase SufS and Its Dynamic Interaction with Frataxin and Scaffold Protein SufU. PLoS One 11, e0158749./li>

30. Kaiser, J. T., Clausen, T., Bourenkow, G. P., Bartunik, H. D., Steinbacher, S., and Huber, R. (2000) Crystal structure of a NifS-like protein from Thermotoga maritima: implications for iron sulphur cluster assembly. J Mol Biol 297, 451–464

31. Fujii, T., Maeda, M., Mihara, H., Kurihara, T., Esaki, N., and Hata, Y. (2000) Structure of a NifS homologue: X-ray structure analysis of CsdB, an Escherichia coli counterpart of mammalian selenocysteine lyase. Biochemistry 39, 1263–1273

32. Lima, C. D. (2002) Analysis of the E. coli NifS CsdB protein at 2.0 A reveals the structural basis for perselenide and persulfide intermediate formation. J Mol Biol 315, 1199–1208

33. Marinoni, E. N., de Oliveira, J. S., Nicolet, Y., Raulfs, E. C., Amara, P., Dean, D. R., and Fontecilla-Camps, J. C. (2012) (IscS-IscU)2 complex structures provide insights into Fe2S2 biogenesis and transfer. Angew Chem Int Ed Engl 51, 5439–5442

34. Shi, R., Proteau, A., Villarroya, M., Moukadiri, I., Zhang, L., Trempe, J. F., Matte, A., Armengod, M. E., and Cygler, M. (2010) Structural basis for Fe-S cluster assembly and tRNA thiolation mediated by IscS protein-protein interactions. PLoS Biol 8, e1000354./li>

35. Muhlenhoff, U., Richhardt, N., Ristow, M., Kispal, G., and Lill, R. (2002) The yeast frataxin homolog Yfh1p plays a specific role in the maturation of cellular Fe/S proteins. Hum Mol Genet 11, 2025–2036

36. Stehling, O., Elsasser, H. P., Bruckel, B., Muhlenhoff, U., and Lill, R. (2004) Iron-sulfur protein maturation in human cells: evidence for a function of frataxin. Hum Mol Genet 13, 3007–3015

37. Ast, T., Meisel, J. D., Patra, S., Wang, H., Grange, R. M. H., Kim, S. H., Calvo, S. E., Orefice, L. L., Nagashima, F., Ichinose, F., Zapol, W. M., Ruvkun, G., Barondeau, D. P., and Mootha, V. K. (2019) Hypoxia Rescues Frataxin Loss by Restoring Iron Sulfur Cluster Biogenesis. Cell 177, 1507–1521 e1516

38. Gervason, S., Larkem, D., Mansour, A. B., Botzanowski, T., Muller, C. S., Pecqueur, L., Le Pavec, G., Delaunay-Moisan, A., Brun, O., Agramunt, J., Grandas, A., Fontecave, M., Schunemann, V., Cianferani, S., Sizun, C., Toledano, M. B., and D’Autreaux, B. (2019) Physiologically relevant reconstitution of iron-sulfur cluster biosynthesis uncovers persulfide-processing functions of ferredoxin-2 and frataxin. Nat Commun 10, p3566

39. Lill, R., and Freibert, S. A. (2020) Mechanisms of Mitochondrial Iron-Sulfur Protein Biogenesis. Annu Rev Biochem 89, 471–499

40. Cai, K., Frederick, R. O., Dashti, H., and Markley, J. L. (2018) Architectural Features of Human Mitochondrial Cysteine Desulfurase Complexes from Crosslinking Mass Spectrometry and Small-Angle X-Ray Scattering. Structure 26, 1127–1136 e1124

41. Cory, S. A., Lin, C. W., Patra, S., Havens, S. M., Putnam, C. D., Russell, D. H., and Barondeau, D. P. (manuscript submitted) Architecture-swapping of the human Fe-S cluster biosynthetic subcomplex: a new morpheein.

42. Maio, N., Jain, A., and Rouault, T. A. (2020) Mammalian iron-sulfur cluster biogenesis: Recent insights into the roles of frataxin, acyl carrier protein and ATPase-mediated transfer to recipient proteins. Curr Opin Chem Biol 55, 34–44

43. Fox, N. G., Martelli, A., Nabhan, J. F., Janz, J., Borkowska, O., Bulawa, C., and Yue, W. W. (2018) Zinc(II) binding on human wild-type ISCU and Met140 variants modulates NFS1 desulfurase activity. Biochimie 152, 211–218

44. Lim, S. C., Friemel, M., Marum, J. E., Tucker, E. J., Bruno, D. L., Riley, L. G., Christodoulou, J., Kirk, E. P., Boneh, A., DeGennaro, C. M., Springer, M., Mootha, V. K., Rouault, T. A., Leimkuhler, S., Thorburn, D. R., and Compton, A.G. (2013) Mutations in LYRM4, encoding iron-sulfur cluster biogenesis factor ISD11, cause deficiency of multiple respiratory chain complexes. Hum Mol Genet 22, 4460–4473

45. Das, D., Patra, S., Bridwell-Rabb, J., and Barondeau, D. P. (2019) Mechanism of frataxin “bypass” in human iron-sulfur cluster biosynthesis with implications for Friedreich’s ataxia. J Biol Chem 294, 9276–9284

46. Behshad, E., and Bollinger, J. M., Jr. (2009) Kinetic analysis of cysteine desulfurase CD0387 from Synechocystis sp. PCC 6803: formation of the persulfide intermediate. Biochemistry 48, 12014–12023

47. Zheng, L., White, R. H., Cash, V. L., and Dean, D. R. (1994) Mechanism for the desulfurization of L-cysteine catalyzed by the nifS gene product. Biochemistry 33, 4714–4720

48. Degli Esposti, M., Mentel, M., Martin, W., and Sousa, F. L. (2019) Oxygen Reductases in Alphaproteobacterial Genomes: Physiological Evolution From Low to High Oxygen Environments. Front Microbiol 10, p499

49. Ball, S. G., Bhattacharya, D., and Weber, A. P. (2016) EVOLUTION. Pathogen to powerhouse. Science 351, 659–660

50. Peterson, E. A., and Sober, H. A. (1954) Preparation of Crystalline Phosphorylated Derivatives of Vitamin B6. Journal of the American Chemical Society 76, 169–175

51. Crooks, G. E., Hon, G., Chandonia, J. M., and Brenner, S. E. (2004) WebLogo: a sequence logo generator. Genome Res 14, 1188–1190

52. Sievers, F., and Higgins, D. G. (2014) Clustal Omega, accurate alignment of very large numbers of sequences. Methods Mol Biol 1079, 105–116

53. Siegel, L. M. (1965) A Direct Microdetermination for Sulfide. Anal Biochem 11, 126–132

54. Margosiak, S. A., Vanderpool, D. L., Sisson, W., Pinko, C., and Kan, C. C. (1996) Dimerization of the human cytomegalovirus protease: kinetic and biochemical characterization of the catalytic homodimer. Biochemistry 35, 5300–5307

55. Laganowsky, A., Reading, E., Allison, T. M., Ulmschneider, M. B., Degiacomi, M. T., Baldwin, A. J., and Robinson, C. V. (2014) Membrane proteins bind lipids selectively to modulate their structure and function. Nature 510, 172–175

