## Supporting Information for "Recapitulating the frataxin activation mechanism in an engineered bacterial cysteine desulfurase supports the architectural switch model"

### Supplementary Information

Department of Chemistry, Texas A&M University, College Station, TX 77842, USA.

**Table S1. Catalytic and binding of IscS variants at position 10, 247 and 248.**

| Complex | $k_{cat}$ ($min^{-1}$) | $K_M$ (μM) | Hill coefficient | $A_s$ ($min^{-1}$) | $K_d$ (μM) |
|---|---|---|---|---|---|
| IscS | 8.2 ± 0.1 | 20 ± 2 | 1 | 18.7 ± 0.1 | 0.008 ± 0.003 |
| $IscS^{S10Q}$ | 1.90 ± 0.05 | 11 ± 2 | 1 | 9.4 ± 0.7 | 1.0 ± 0.6 |
| $IscS^{PL}$ | 4.12 ± 0.07 | 18 ± 2 | 1 | 18.8 ± 0.8 | 1.4 ± 0.3 |
| $IscS^{QPL}$ | 0.22 ± 0.01 | 39 ± 9 | 1 | 5.8 ± 0.5 | 32 ± 7 |
| $IscS^{NPL}$ | 8.86 ± 0.12 | 24 ± 1 | 1.39 ± 0.08 | 28 ± 2 | 0.8 ± 0.3 |
| $IscS^{HPL}$ | 1.51 ± 0.05 | 37 ± 4 | 1.6 ± 0.2 | 14.2 ± 0.8 | 8 ± 1 |

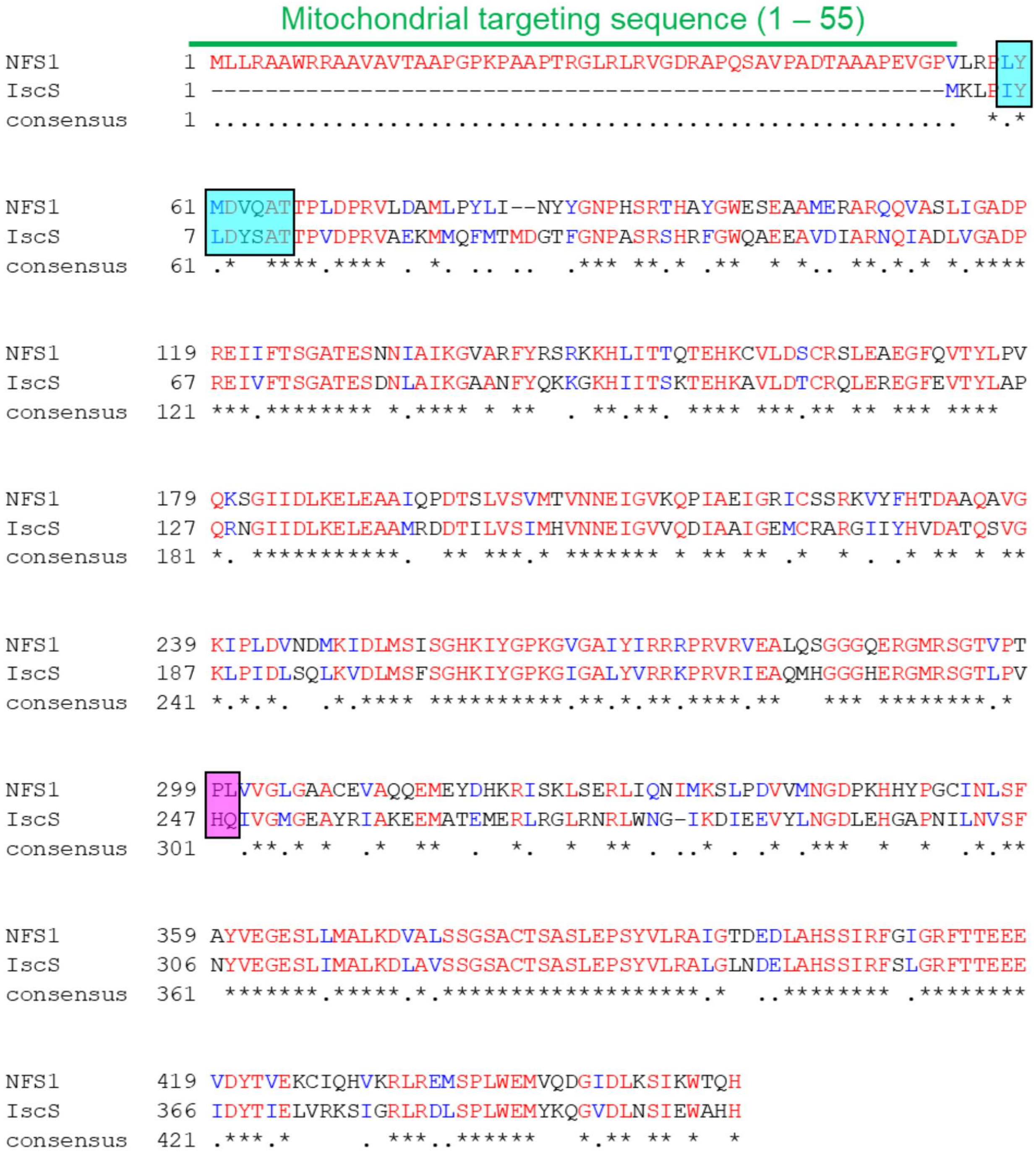


**Figure S1**. **Sequence alignment of human NFS1 and *E. coli* IscS.** The sequences were aligned using Clustal Omega default parameters and displayed with BoxShade. The mitochondrial targeting sequence (residues 1-55) are displayed in green whereas residues L59-T66 are displayed using an aqua rectangle. The residues P299 and L300 are highlighted with a purple rectangle. Residues in red with stars are identical and those in blue are conservative substitutions.

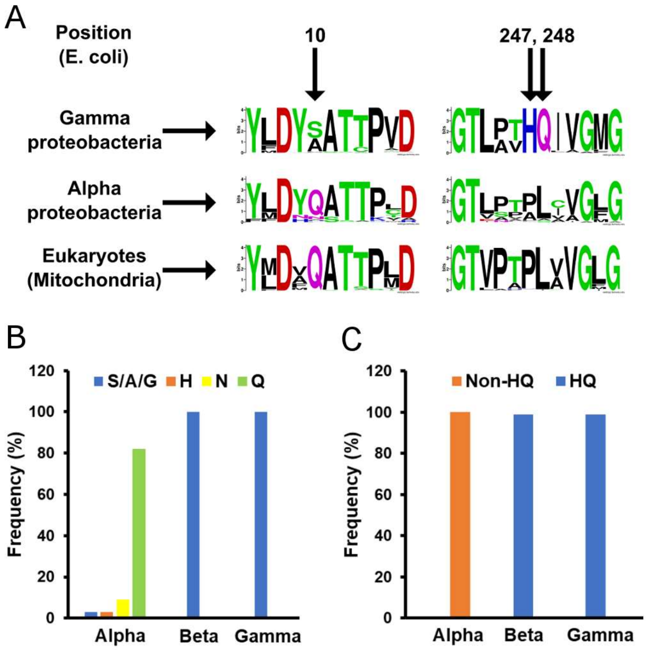


**Figure S2. Sequence motif conservation in proteobacteria and eukaryotic cysteine desulfurases.** Homologous sequences to human NFS1 were identified in proteobacteria and eukaryotes, manually curated, and aligned with Clustal Omega. (**A**) Sequence logo were generated with WebLogo for cysteine desulfurase motifs from γ-proteobacteria (*E. coli* is a member), α-proteobacteria, and eukaryotic mitochondria. The frequency of residue corresponding to (**B**) position 10 and (**C**) the HQ pair (H247, Q248) from *E. coli* IscS are displayed for different proteobacteria. The S/A/G refers to either a serine, alanine, or glycine residue. Motifs other than HQ are indicated by the non-HQ designation.

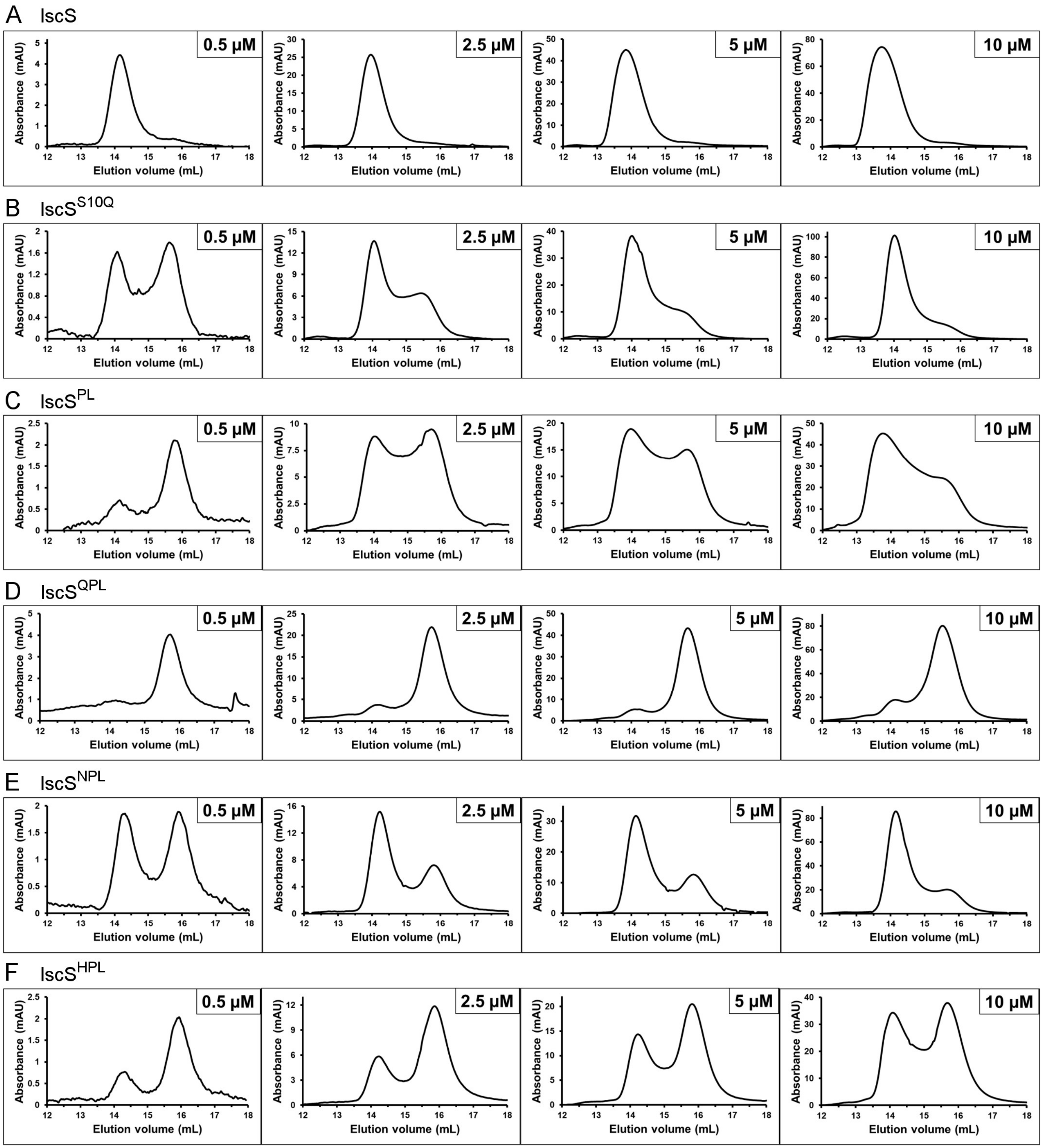


**Figure S3. Analysis of oligomeric state of IscS variants with size exclusion chromatography.** Protein samples (0.5, 2.5, 5 and 10 µM) were analyzed anaerobically using an analytical S200 column in the presence of 2.5 mM TCEP. The peaks near 14 mL and 15.8 mL correlate with the dimeric and monomeric forms of IscS, respectively. Samples were analyzed for (**A**) native IscS and the (**B**) IscS$^{S10Q}$, (**C**) IscS$^{PL}$, (**D**) IscS$^{QPL}$, (**E**) IscS$^{NPL}$, and (**F**) IscS$^{HPL}$ variants. The increased dimer population observed for IscS variants at higher protein concentrations is consistent with an equilibrium between monomeric and dimeric species.

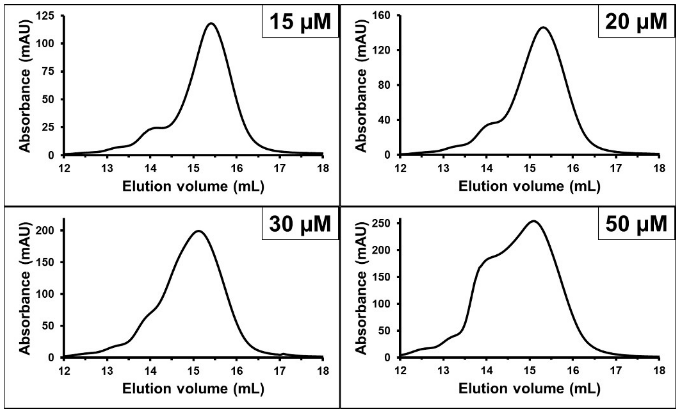


**Figure S4. Analysis of oligomeric state of the IscS$^{QPL}$ variant at higher protein concentrations with SEC.** The quaternary structure of the IscS$^{QPL}$ variant was analyzed at high concentrations (15, 20, 30 and 50 μM) with SEC conducted under anaerobic conditions in the presence of 2.5 mM TCEP. The peaks near 14 mL and 15.8 mL correlate with the dimeric and monomeric forms of IscS, respectively.

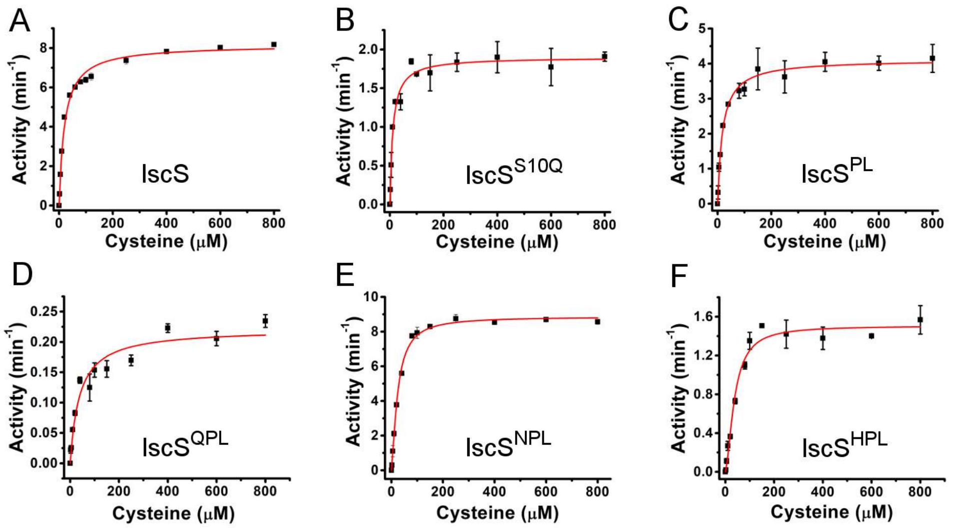


**Figure S5. The cysteine desulfurase activity for native IscS and IscS variants.** (**A**) Native IscS and the (**B**) $IscS^{S10Q}$, (**C**) $IscS^{PL}$, (**D**) $IscS^{QPL}$, (**E**) $IscS^{NPL}$, and (**F**) $IscS^{HPL}$ variants (0.5 μM) were anaerobically reacted with different concentrations of L-cysteine in the presence of 4 mM D,L-DTT at 37 °C for 6 min. The sulfide produced was converted to methylene blue using a DPD and $FeCl_3$ mixture and quantified using a sulfide standard curve. The data is fit with a Michaelis-Menten equation that includes cooperativity. Error bars represent replicate trials (n = 3).

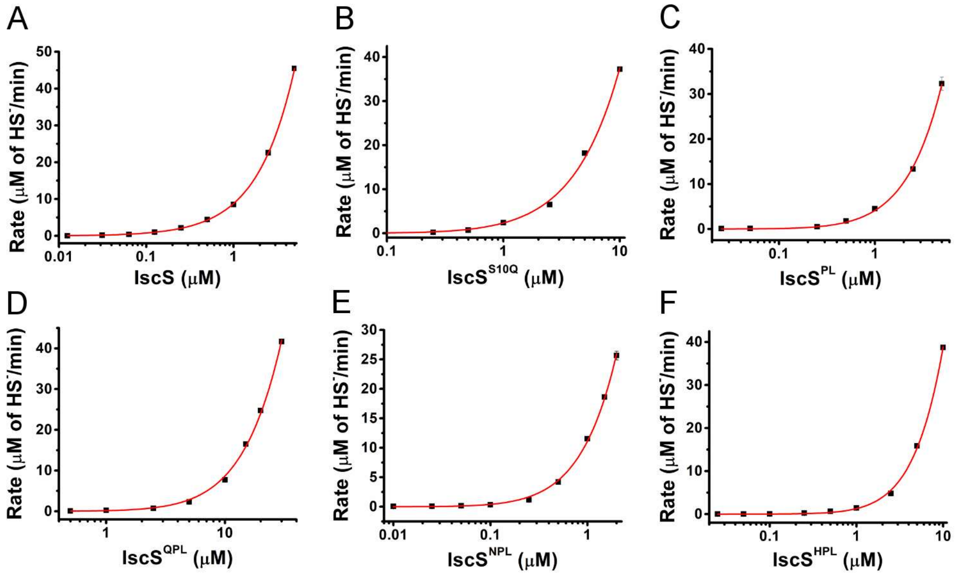


**Figure S6. The S10Q, H247P and Q248L substitutions reduce the affinity between the cysteine desulfurase subunits.** Cysteine desulfurase activities at different concentrations of (**A**) native IscS and the (**B**) $IscS^{S10Q}$, (**C**) $IscS^{PL}$, (**D**) $IscS^{QPL}$, (**E**) $IscS^{NPL}$, and (**F**) $IscS^{HPL}$ variants were determined using 1 mM of L-cysteine in the presence of 4 mM D,L-DTT at 37 °C. The rates of sulfide generation were determined as before (see Fig. S5), plotted against the concentration of enzyme, and fitted (red line; see Methods) to determine the dimer dissociation constants and dimeric activities.

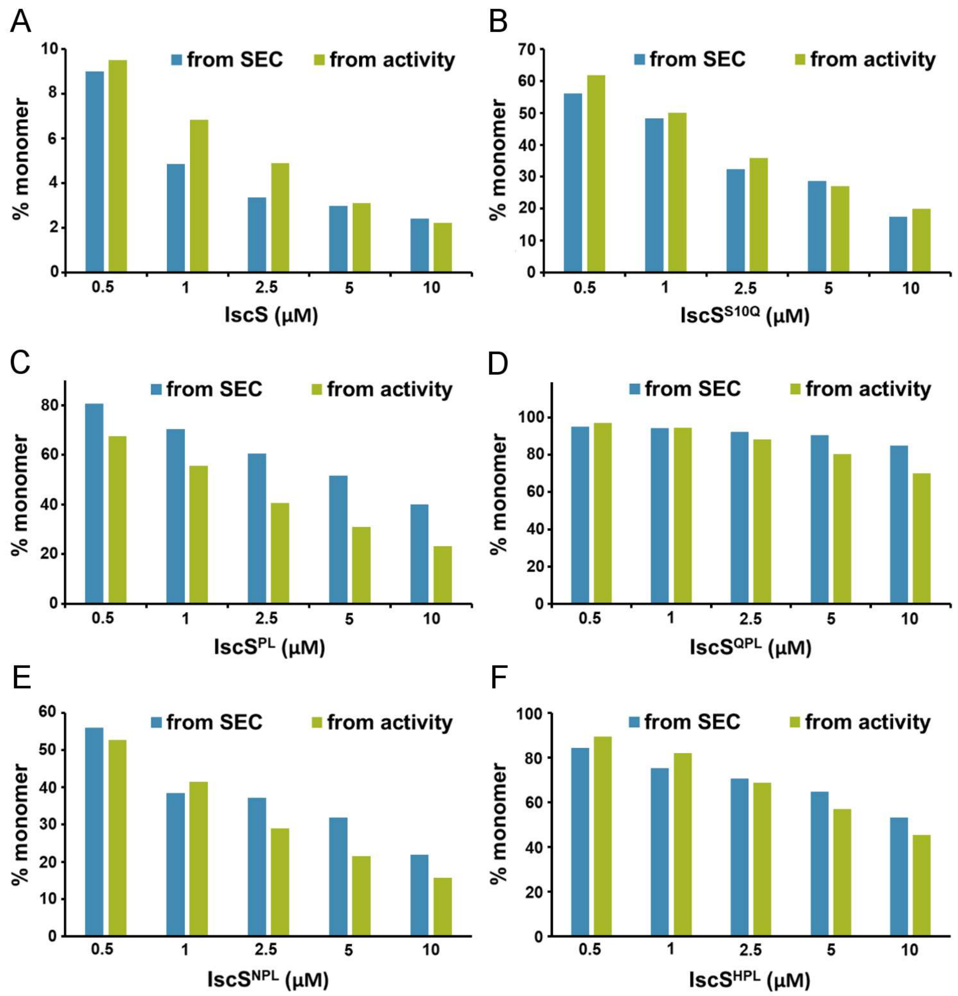


**Figure S7. Comparison of the percent monomer calculated from dimer dissociation constants and analytical SEC data.** The percent monomer from SEC analysis (blue) and activity (green) displayed for (**A**) native IscS and the (**B**) IscS$^{S10Q}$, (**C**) IscS$^{PL}$, (**D**) IscS$^{QPL}$, (**E**) IscS$^{NPL}$, and (**F**) IscS$^{HPL}$ variants. The dimer dissociation constants were used to calculate the percent monomer for each of the variants at the different protein concentrations. The percent monomer was also estimated by dividing the area under the monomer peak (~15.4 – 17 mL elution volume) from analytical SEC experiments with the total area (monomer + dimer) followed by multiplication by 100.

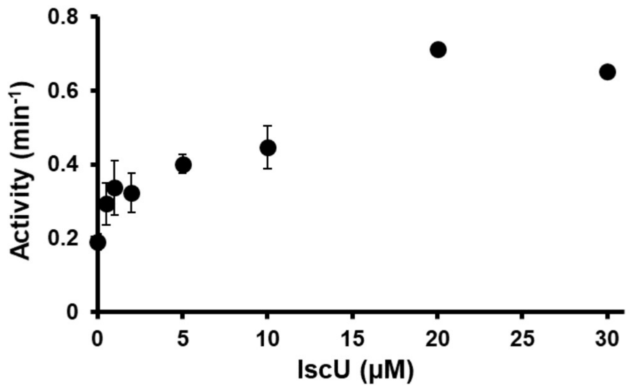


**Figure S8. IscU enhances activity of the IscS$^{QPL}$ variant.** The cysteine desulfurase activity was determined using 0.5 μM of the IscS$^{QPL}$ variant, 1 mM L-cysteine and 4 mM D, L-DTT at 37 °C under anaerobic condition with varying amounts of IscU (0 – 30 μM). The activity increased with increasing amounts of IscU and saturated at about 20 μM. Error bars are replicate errors (n =3).

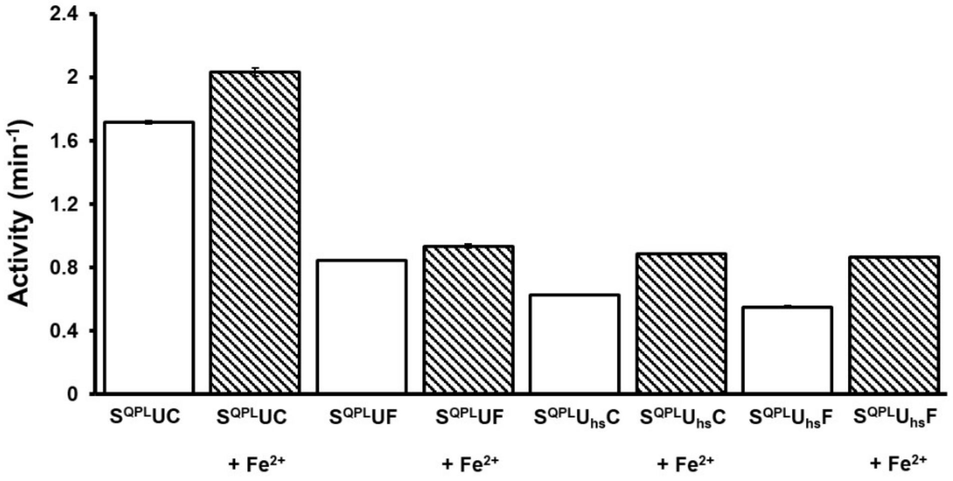


**Figure S9. Ferrous iron further enhances the cysteine desulfurase activity of the IscS$^{QPL}$ variant in the presence of the scaffold protein and CyaY/FXN.** The activity was determined using 0.5 μM of the IscS$^{QPL}$ variant, 30 μM of IscU/ISCU2, 20 μM of CyaY/FXN, 1 mM L-cysteine, 10 μM $Fe^{2+}$ and 4 mM D, L-DTT at 37 °C under anaerobic condition. Error bars represent replicate errors (n = 3). Abbreviations: IscS$^{QPL}$ = S$^{QPL}$, IscU = U, ISCU2 = U$_{hs}$, FXN = F, CyaY = C.

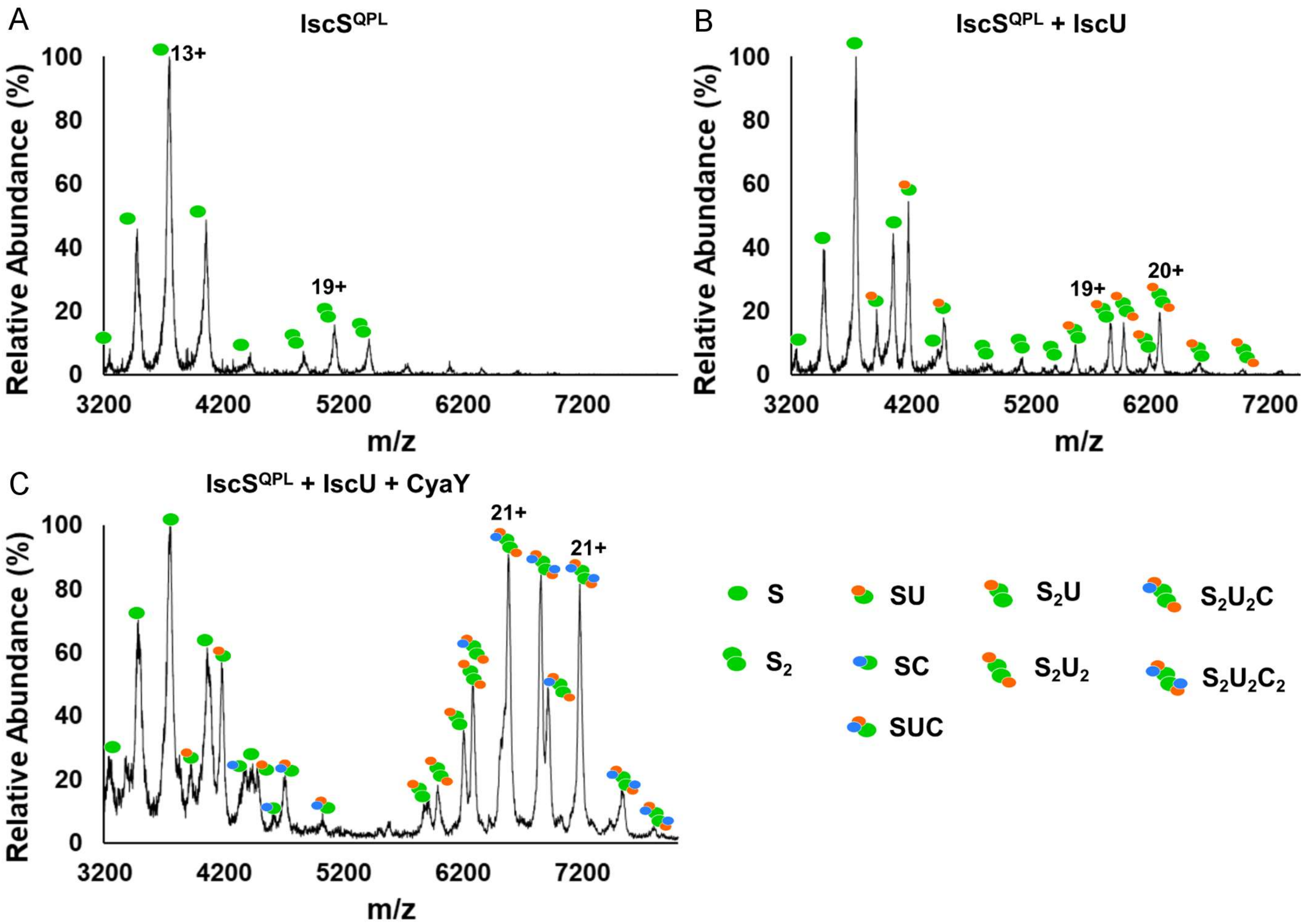


**Figure S10. Native MS spectra for the IscS$^{QPL}$ variant in complex with IscU and with both IscU and CyaY.** The native MS spectra are displayed for the **(A)** 4 μM IscS$^{QPL}$ variant, **(B)** 4 μM IscS$^{QPL}$ variant with 6 μM IscU, and (**C)** 4 μM IscS$^{QPL}$ variant with 6 μM IscU and 12 μM CyaY. Some of the charge states are indicated. The compositions of the different complexes are shown above the MS peaks and defined on the right side of the figure. Abbreviations IscS$^{QPL}$ variant = S; IscU = U; CyaY = C.

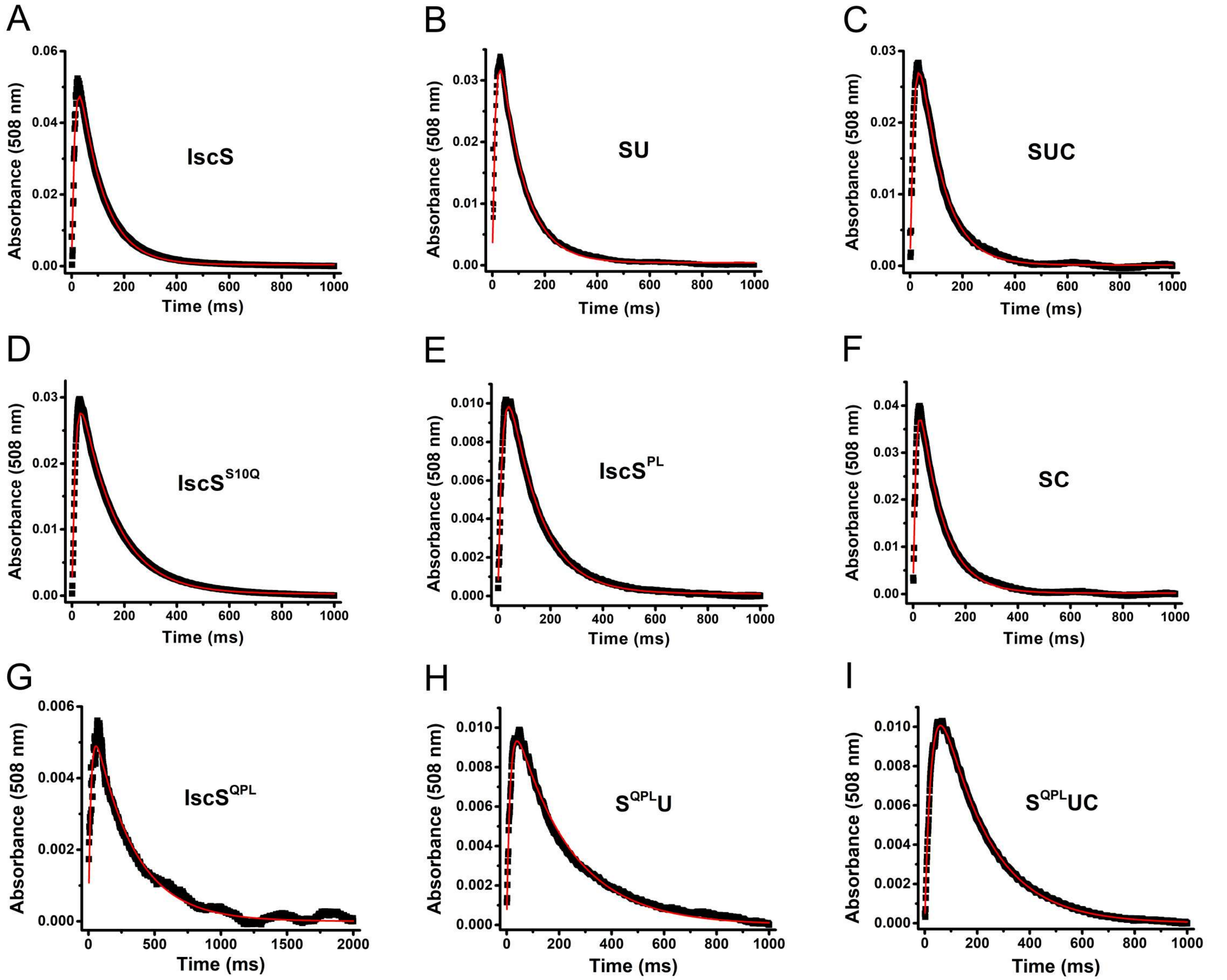


**Figure S11. Formation and decay kinetics of the Cys-quinonoid intermediate for IscS variants and complexes.** The quinonoid decay kinetics were measured using stopped-flow spectroscopy. 30 μM (final concentration) of IscS or IscS variants alone or in combination with IscU (30 μM) and/or CyaY (30 μM) were mixed with 5 mM L-cysteine (final concentration). The formation and decay of the Cys-quinonoid intermediate was followed by the absorbance change at 508 nm for 1 minute. The rates for the formation and decay of the (**A**) IscS, (**B**) IscS-IscU (SU), (**C**) IscS-IscU-CyaY (SUC), (**D**) IscS$^{S10Q}$, (**E**) IscS$^{PL}$, (**F**) IscS-CyaY (SC), (**G**) IscS$^{QPL}$, (**H**) IscS$^{QPL}$-IscU (S$^{QPL}$U), and (**I**) IscS$^{QPL}$-IscU-CyaY (S$^{QPL}$UC) complexes were obtained by fitting (red traces) the change in absorbance at 508 nm with time. The decay kinetics are tabulated and compared in Figure. 7.

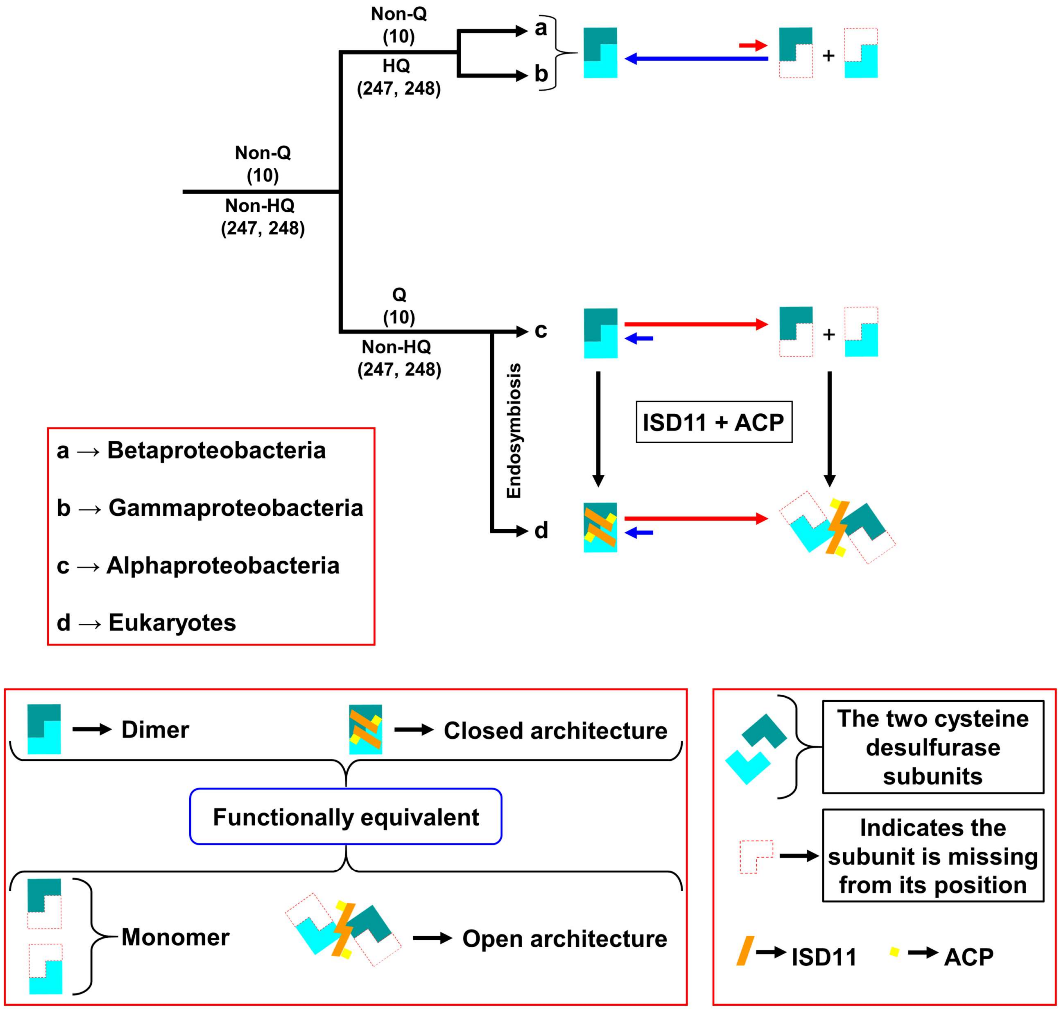


**Figure S12. Model depicting the evolution of the eukaryotic cysteine desulfurases.** The cysteine desulfurase of the last common ancestor (LCA) of the α-, β- and γ-proteobacteria most likely had a non-Q residue (any of S, A, G, H, N but not Q) at position 10 and non-HQ residues at the equivalent 247 and 248 positions of the *E. coli* enzyme. Since only the presence of a Q at position 10 and absence of the HQ pair leads to a substantially weaker dimer interface, the LCA cysteine desulfurase subunits most likely had a moderate affinity and existed as a mixture of monomeric and dimeric species. During evolution, both β- and γ-proteobacteria retained the non-Q residue but acquired the HQ pair. Consequently, the cysteine desulfurase in these clades likely exist as primarily dimeric enzymes. On the other hand, the α-proteobacteria retained the non-HQ residue while acquiring a glutamine at position 10. As a result, these cysteine desulfurases likely exist as primarily monomeric species with a weak dimer interface. After the endosymbiosis, at some point during eukaryogenesis, the cysteine desulfurase acquired the eukaryotic specific ISD11, which interacts with the already present mitochondrial ACP. The addition of these subunits converted the dimeric cysteine desulfurase into the ready architecture and facilitated the formation of the open architecture from the monomeric cysteine desulfurase. Since the dimer interface remained very weak, the open form became the predominant architecture.

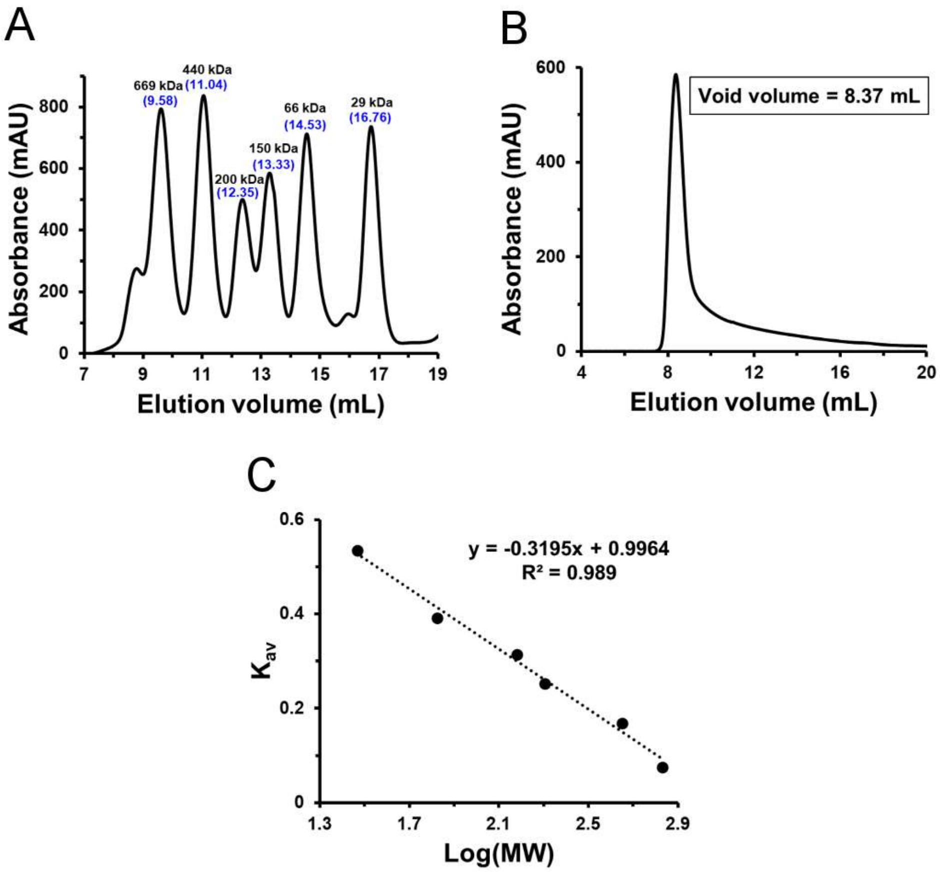


**Figure S13. Standard curve for analytical size exclusion chromatography. (A)** Chromatogram for molecular weight standards. Molecular weights and elution volumes are given in black and blue (within parenthesis), respectively. **(B)** Blue dextran (MW ~ $10^6$) was used to determine a void volume of 8.37 mL. **(C)** $K_{av}$ was calculated using the elution volume of the standards, void volume, and column volume as described in the Methods. $K_{av}$ was plotted against the log of the molecular weight (MW) and fitted to a linear equation.
